# The 5:2 diet does not increase adult hippocampal neurogenesis or enhance spatial memory in mice

**DOI:** 10.1101/2022.10.03.510613

**Authors:** Luke D. Roberts, Amanda K. E. Hornsby, Alanna Thomas, Martina Sassi, Aimee Kinzett, Nathan Hsiao, Bethan R David, Mark Good, Timothy Wells, Jeffrey S. Davies

**Author notes:** Equal contribution (Luke D. Roberts, Amanda K. E. Hornsby).

## Abstract

New neurones are generated throughout life in the mammalian brain in a process known as adult hippocampal neurogenesis (AHN). Since this phenomenon grants a high degree of neuroplasticity influencing learning and memory and mood related behaviour, identifying factors that regulate AHN may be important for ameliorating age-related cognitive decline and neurodegeneration. Calorie restriction (CR), in the absence of malnutrition, has been shown to enhance AHN and improve hippocampal-dependent memory, mediated by the stomach hormone, ghrelin. Intermittent fasting (IF), a dietary strategy offering more flexibility than conventional CR, also promotes aspects of AHN. The 5:2 diet is a popular form of IF linked to a range of health benefits, however its effects on AHN and spatial memory are not well characterised. We hypothesised that the 5:2 diet would enhance AHN in a ghrelin-dependent manner.

To assess this, we used immunohistochemistry to quantify new adult-born neurones and new neural stem cells (NSCs) in the hippocampal DG of adolescent and adult wild-type and mice lacking the ghrelin receptor following six weeks on a 5:2 diet. We report an age-related decline in neurogenic processes and identify a novel role for ghrelin-receptor in regulating the formation of new adult born NSCs in an age-dependent manner. However, the 5:2 diet did not affect new neurone or NSC formation in the DG. Consistent with this finding the 5:2 diet did not alter performance on a spatial learning and memory task. These data suggest that the 5:2 diet used in this study does not increase AHN or improve associated spatial memory function.

**Highlights:**

- 5:2 diet does not increase adult hippocampal neurogenesis
- 5:2 diet does not enhance spatial memory performance
- There is an age-related decline in adult hippocampal neurogenesis
- Ghrelin-receptor regulates new neural stem cell and new neurone number in an age-related manner
- Ghrelin-receptor is required for intact spatial memory

## Introduction

In order to effectively combat age-related morbidities, both in the brain and at the systemic level, appropriate interventions that target the underlying molecular hallmarks are required. A promising candidate in this regard is dietary restriction (DR), a group of interventions that reduce either daily caloric intake, macronutrient intake, feeding duration or a mixture of each, without inducing malnutrition (Di Francesco et al., 2018; Fontana & Partridge 2015). These paradigms have emerged as robust interventions for delaying and preventing the incidence of age-related disease and metabolic dysfunction in rodents and non-human primates (Di Francesco et al., 2018; Fontana & Partridge, 2015; Mattison et al., 2017).

While DR exists in numerous forms, notable examples include calorie restriction (CR), where daily calorie intake is reduced (typically by 20-40%) but meal frequency and timing is not constrained; and intermittent fasting (IF) diets, where individuals undergo extended periods (16-48h) with very limited or no food intake, with the resumption of *ad-libitum* eating habits in between (Di Francesco et al., 2018; Mattson et al., 2017).

There are likely to be myriad cellular and molecular mechanisms underlying the beneficial effects of DR (Di Francesco et al., 2018; Mattson et al., 2018). However, a notable paradigm implicated in DR regulated brain plasticity is the birth of new neurones in the adult hippocampus (Buntwal et al., 2019; Morgan et al., 2017). This phenomenon, known as adult hippocampal neurogenesis (AHN), provides a high degree of neuroplasticity and regulates spatial memory function and mood-like behaviours. AHN declines markedly with age, therefore, identifying factors that regulate AHN are potentially significant for attenuating age-related cognitive decline (Mu & Gage, 2011).

IF and periodic fasting mimicking diets (FMD) (Brandhorst et al., 2015) have been shown to promote aspects of neurogenesis including cell proliferation (Brandhorst et al.,2015; Dias et al., 2021), new cell survival (Dias et al., 2021; Lee et al., 2000; Lee et al., 2002a; Lee et al., 2002b) and neurone differentiation (Dias et al., 2021). Moreover, CR increases neurone differentiation, the number of new adult born hippocampal neurones and improves hippocampal-dependent cognition by signalling via the stomach hormone, ghrelin (Hornsby et al., 2016).

Ghrelin, which exists as acylated and unacylated forms in the circulation, is secreted from the stomach during periods of fasting (Tschöp *et al*, 2000) to stimulate feeding (Nakazato *et al*, 2001) and glucose homesostasis (Mani & Zigman, 2017). Exogenous administration of acyl-ghrelin stimulated the proliferation and neuronal differentiation of hippocampal progenitors in adult mice (Moon et al. 2009). Subsequently, cell proliferation in the DG, as well as the proportion of new BrdU^+^ cells that express markers of immature neurones (BrdU^+^/DCX^+^), neurone differentiation (% of BrdU^+^/NeuN^+^) and mature neurones (BrdU^+^/NeuN^+^), were reported to be decreased in the ghrelin-knockout (GKO) mice (Li et al., 2013). Ghrelin administration restored levels of neuronal differentiation to those of WT controls, supporting its pro-neurogenic role (Li et al., 2013). In addition, the stress-induced reduction in AHN in a mouse model of chronic psychosocial stress is more severe in mice lacking the ghrelin-receptor, growth hormone secretagogue receptor (GHS-R) (Walker et al., 2015).

Notably, we showed that two-weeks of daily acyl-ghrelin injections, at physiological levels, significantly increased the number of new adult-born neurones in the dentate gyrus of the hippocampus. The acyl-ghrelin-treated rats also displayed enhanced pattern separation performance in a Spontaneous Location Recognition task conducted 8-10 days following the end of treatment, indicating that acyl-ghrelin had a long-lasting effect on spatial memory (Kent et al., 2015). Similarly, our data suggested that the beneficial effects of CR on AHN are also mediated by acyl-ghrelin as a short period of 30% CR increased the number of new adult-born hippocampal neurones in a ghrelin-receptor dependent manner (Hornsby et al., 2016). Wild-type and GKO mice maintained for 3 months on alternate day fasting (ADF) or *ad libitum* diet reported that DR increased the number of new adult-born cells in wild-type mice, but not in GKO mice, suggesting that acyl-ghrelin signalling is required for the beneficial effects of DR on aspects of neurogenesis (Kim et al., 2015). Daily systemic injections of acyl-ghrelin for 8 days enhanced AHN (Zhao et al.2014) and acyl-ghrelin treatment rescued abnormal neurogenesis in a mouse model of AD (5xFAD) (Moon et al., 2014). These studies provide strong evidence in support of the pro-neurogenic role of acyl-ghrelin signalling (Kent et al., 2015).

The precise mechanisms by which acyl-ghrelin signalling mediates AHN remains unclear, however, this may be via non-cell autonomous pathways involving the release of pro-neurogenic factors (Buntwal et al., 2019; Sassi et al., 2022). More recently, we showed that acyl-ghrelin increased BDNF mRNA – a neurotrophic factor - within the granule cell layer of the hippocampus (Hornsby et al.2020). Consistent with an increase in BDNF, acyl-ghrelin binding to GHS-R on hippocampal neurones promoted dendritic spine formation, increased synaptic plasticity and LTP (Diano et al., 2006).

The 5:2 diet is a popular form of IF in which the usual calorie intake is restricted on 2 non-consecutive days within the week, while a normal diet is consumed on the remaining 5 days (Patterson et al., 2015). Reducing the normal calorie intake to 25% for 2 days per week improves several health biomarkers (Harvie et al., 2011). However, unlike for conventional CR and ADF diets, studies exploring the impact of the 5:2 diet on brain health and AHN are limited. One study reported that a four-week 5:2 diet improved pattern separation performance but impaired memory retention in an adult population with central obesity (Kim et al., 2020). As relatively little is known about the impact of the 5:2 diet on AHN and pattern separation memory, we address this gap in knowledge by testing the hypothesis that the 5:2 diet promotes AHN and cognition via ghrelin-receptor signalling.

## Results

### The 5:2 diet did not reduce body weight in female mice

To assess the effect of the 5:2 diet on AHN and underlying involvement of ghrelin signalling, adolescent (7-week old) and adult (7-month old) homozygous loxTB-GHSR (herein referred to as GHS-R^−/−^) mice and their wild type (WT) littermate equivalents were subjected to two non-consecutive fasting days (Monday and Thursday) per week for a period of 6 weeks (Fig 1A). To avoid the potential metabolic consequences of disrupting the circadian regulation of food intake (Glad et al.2011), chow was returned to IF mice at 16.00h, 2 hours before lights off, on re-feeding days (Tuesday and Friday). This was followed by a week of ad libitum (AL) feeding, in which hippocampal learning and memory was assessed via the object in place (OIP) task. Control animals were fed AL for the entire duration of the study.

**Figure 1.**
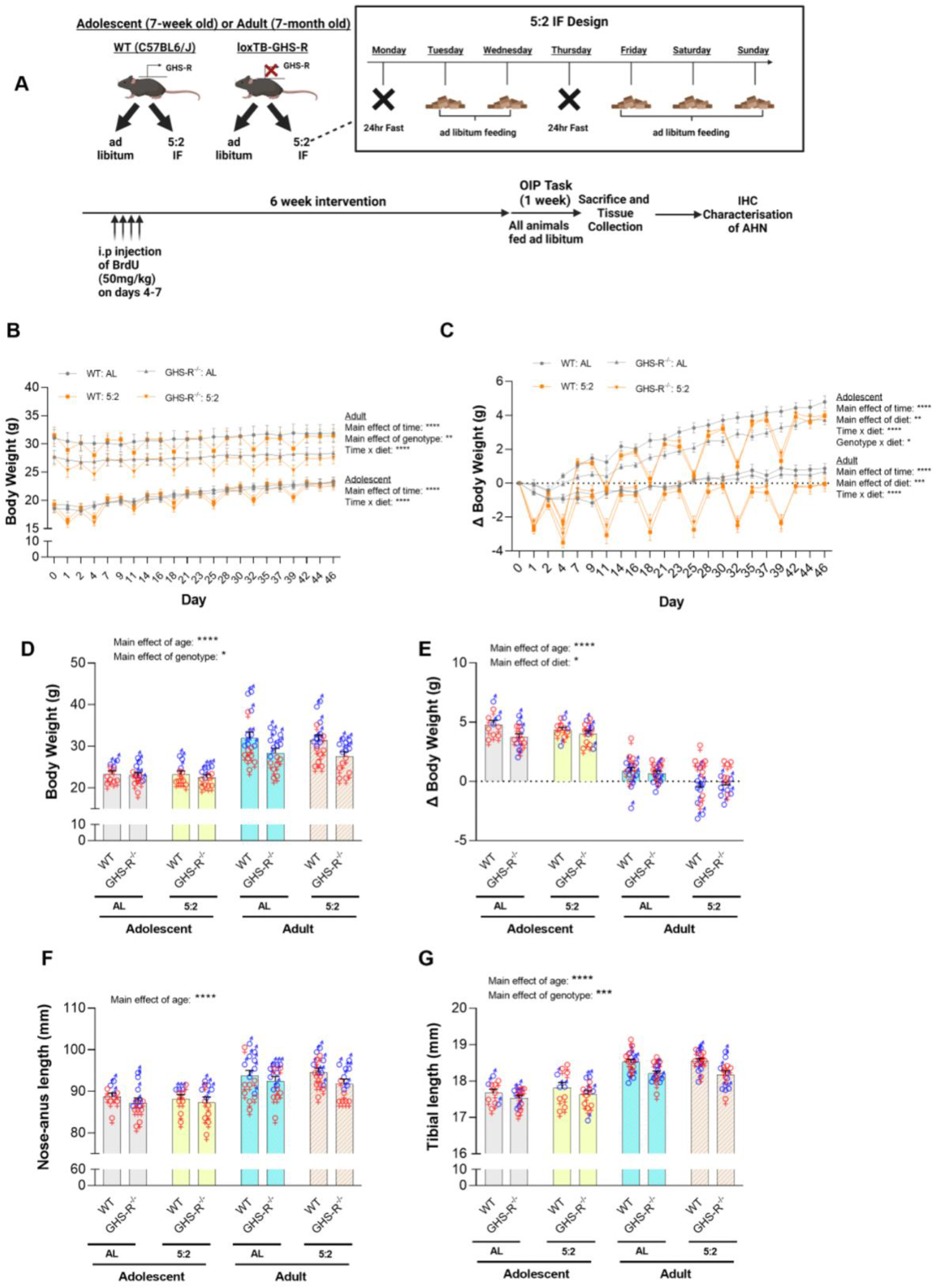
Body weight and growth on the 5:2 diet. Schematic of the experimental timeline (A). Body weight (B) and (C) change in body weight across the course of the study. Final day body weight (D) and change in body weight (E) at the end of the study. Nose-anus (F) and tibial (G) length at the end of the study.

Body weight and growth measures at the end of the study indicated that adult animals had significantly greater body weight (Fig 1D; P = < 0.0001), nose-anus length (Fig 1E; P = < 0.0001) and tibial length (Fig 1F; P = < 0.0001) compared to adolescent mice, while adolescent mice gained significantly more weight (Fig 1G; P = < 0.0001), compared to adult mice. Moreover, loss of GHS-R expression resulted in significant reductions in both body weight (main effect of genotype: P = 0.0097) and tibial length (main effect of genotype: P = 0.0002). The 5:2 diet did not affect final day body weight (main effect of diet: P = 0.5604), nose-anus length (main effect of diet: P = 0.9620) or tibial length (main effect of diet: P = 0.3709); but did result in a modest reduction (compared to AL controls) in weight change relative to baseline measurements (main effect of diet: P = 0.0173). 5:2 fed animals did display considerable body weight fluctuations during the 6-week intervention, with repeated measures analyses identifying significant interactions between time and diet for both body weight (Fig 1B; P = < 0.0001, adolescents and adults) and body weight change relative to baseline (Fig 1C; P = <0.0001, adolescents and adults). Indeed, 5:2 fed animals lost weight in response to fasting and subsequently regained this weight upon accessing food (Fig 1B and 1C). These 5:2 mediated body weight fluctuations were observed for both age groups and both genotypes and were subsequently normalized following the recommencement of AL feeding during the week of cognitive testing. Thus, the 5:2 diet appears to result in elevated rebound body weight on re-feeding days. Interestingly, when the cohorts were separated by sex, the 5:2 diet resulted in a 31% reduction in body weight gain in adolescent WT males and a 9% reduction in adolescent females, relative to ad-libitum fed mice of the same age (Supplementary Fig 1; Sex × Diet interaction P = 0.0316). Therefore, from a broader metabolic perspective, we show that the 5:2 diet is more effective in restricting weight gain in adolescent male mice.

### Cell proliferation in the hippocampus was not altered by the 5:2 diet

To assess the effect of the 5:2 diet on cell proliferation (including Neural Stem Cells (NSCs)) in the DG of the hippocampus, we quantified Ki67 positive cells using DAB immunohistochemistry (Fig 2A). Our findings indicate that while the number of proliferating cells declined with age (Main effect of age: P = < 0.0001), neither the 5:2 diet (Main effect of diet: P = 0.7122) nor loss of GHS-R expression (Main effect of genotype: P = 0.2852) affected the abundance of proliferating cells (Fig 2B). This is consistent with our previous CR study (Hornsby et al.2016) and ADF studies from other research groups (Lee et al.2000; Lee et al. 2002; Bondolfi et al.2004, Kim et al.2015).

**Figure 2.**
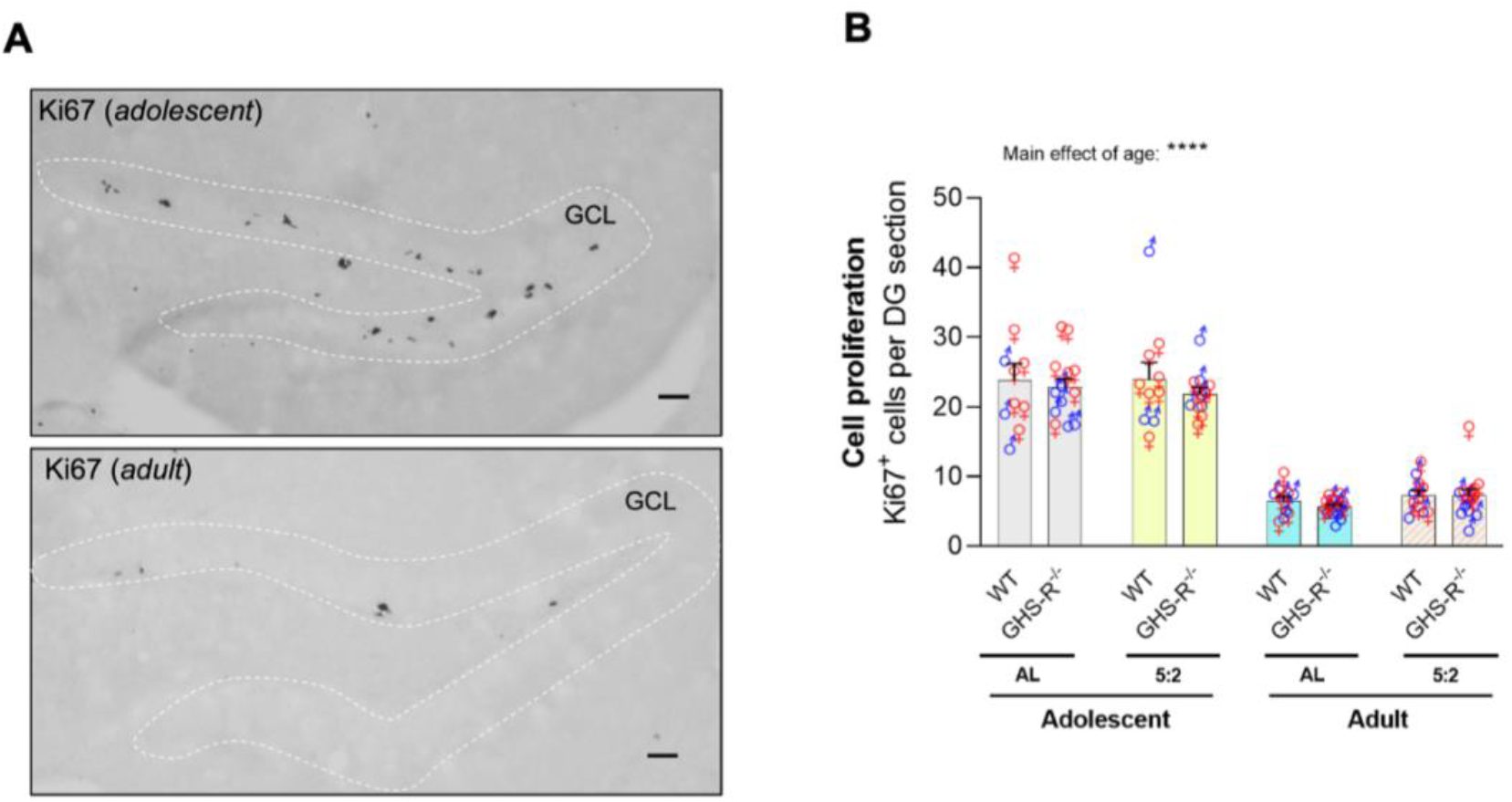
Cell proliferation, new adult born cells and neurones. Representative images of (A) cell division marker, Ki67, in adolescent and adult hippocampus (scale bar = 50um).

### The 5:2 diet increased the number of immature neurones in the hippocampus independently of GHS-R

Next, we quantified immature neurones in the DG using DAB-based IHC against doublecortin (DCX; Fig 3A). We observed a striking age-related decline in the number of DCX positive cells, with adult animals having considerably fewer immature neurones than adolescent animals, regardless of diet or genotype (Fig 3B, main effect of age: P = < 0.0001). Interestingly, a significant main effect of diet was also identified (P = 0.0230) with 5:2 fed animals having more immature neurones compared to those fed AL, and mean differences being more pronounced for adolescent animals. Moreover, loss of GHS-R expression did not attenuate the 5:2 mediated increase in DCX positive cell number, suggesting that the mild effect of the 5:2 diet on increasing the number of immature neurones is independent of ghrelin signalling.

**Figure 3.**
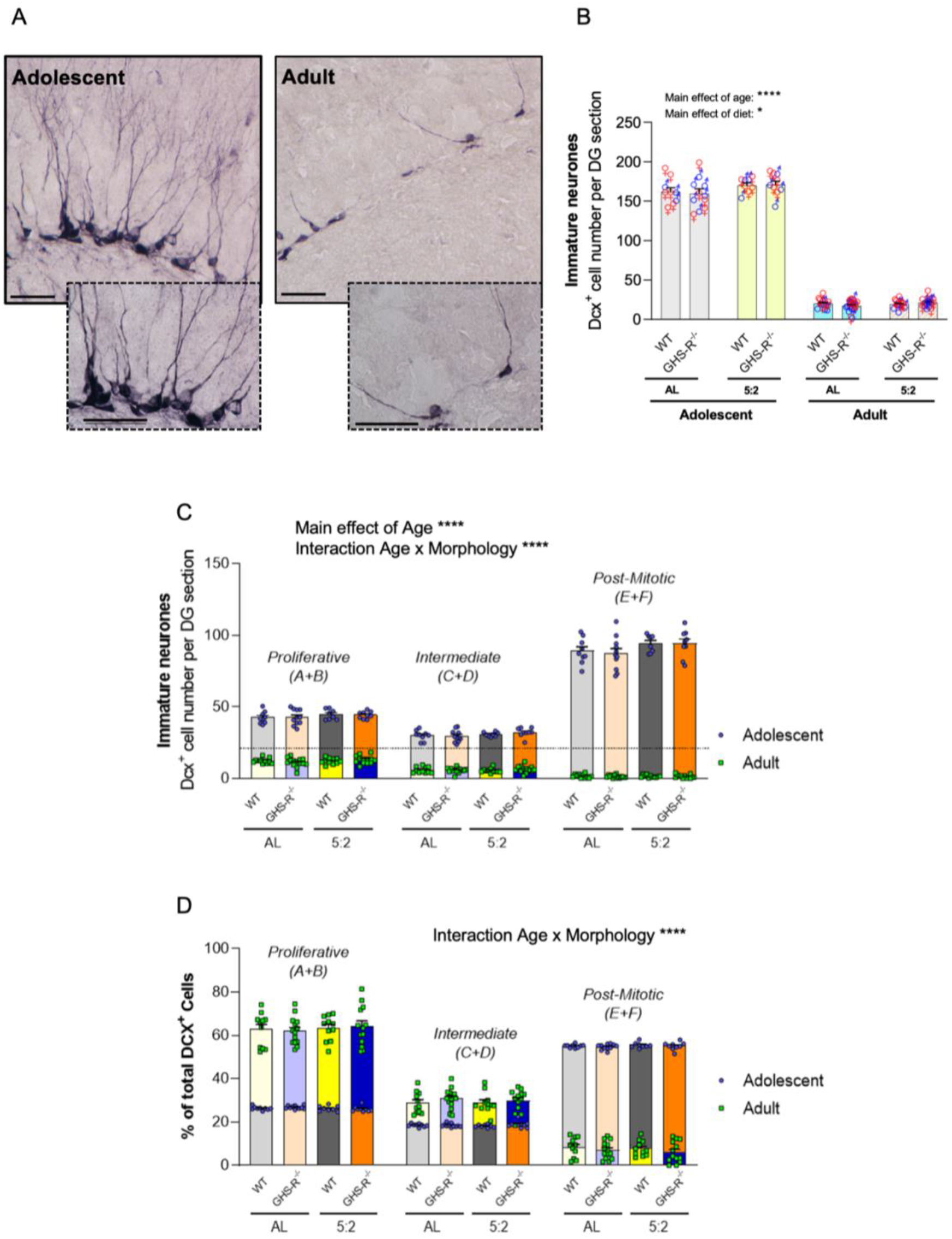
Immature neurone number and morphology in the adult hippocampus. (A) Representative images of Dex immunopositive cells in adolescent and adult mouse hippocampus. Scale bar = 50um. (B) Quantification of hippocampal Dcx^+^ cell number. (C) Quantification of Dcx^+^ cell morphology. (D) Relative abundance of each Dcx^+^ morphological sub-type.

While DCX is generally used as a marker of immature neurones, the period of DCX expression during AHN (~ 3 weeks) covers both proliferative (type 2b NSCs & neuroblasts) and post-mitotic stages (Plümpe et al., 2006). DCX immune-positive cells therefore display a range of morphologies that are representative of distinct phases of neurogenic regulation and developing electrophysiological properties (Plümpe et al., 2006). Thus, in accordance with established criteria (Plümpe et al.2006) we assessed the absolute number (per DG section) (Fig 3C) and relative distributions (% of total) (Fig 3D) of proliferative (A-B), intermediate (C-D), and post-mitotic (E-F) DCX expressing cell morphologies.

As with the total DCX counts, adult animals had fewer DCX positive cells in each morphological category compared to adolescent animals (P = < 0.0001), with the age-related decline being most prominent for post-mitotic cells (Fig 3C). Indeed, comparison of the relative distribution of DCX morphologies revealed an age-related sh ft from a predominantly post-mitotic morphology in adolescents (post-mitotic = ~55% of total DCX cells; proliferative = ~26% of total DCX cells) to a predominantly proliferative morphology in adults (post-mitotic = ~7% of total DCX cells; proliferative = ~63% of total DCX cells) (morphology × age: P = < 0.0001) (Fig 3D). These data suggest that the ageing niche may lack the mechanisms to support maturation of new immature neurones.

No significant main effects of genotype or diet were detected for either adolescent or adult mice, although a significant interaction between morphology and diet was identified for the adolescent age group (P = 0.0387) (Fig 3C). Similarly, a significant interaction between morphology, age and diet was identified when repeated measures 3-way ANOVA was performed on only the GHS-R^−/−^ mice (P = 0.0241). A summary of the statistical analyses can be found in Tables S7-S10.

Our observations that post-mitotic DCX expressing cells pre-dominate over proliferative DCX expressing cells in adolescent mice (7 weeks old at start of study) are consistent with those of Plümpe et al., (2006), who characterised DCX morphologies in 6-week old female mice.

### The number of new adult-born cells in the hippocampus was not altered by the 5:2 diet

Next, we explored the effect of the 5:2 diet on the generation of new adult-born cells in the hippocampal DG in both wild-type and GHS-R^−/−^ mice, from adolescent and adult populations. To do this, a BrdU-pulse chase method was used with all mice receiving twice-daily injections of BrdU on the first two days of the study and BrdU^+^ immunoreactive cells in the SGZ and GCL were counted six weeks later (Fig 4A). The mean number of BrdU^+^ cells was quantified for each mouse and showed there was no effect of diet on the number of BrdU^+^ cells in the SGZ and GCL (main effect P = 0.2948; Fig 4B), suggesting that this six week 5:2 regimen does not increase the generation of new adult-born cells in the DG. In contrast, Dias et al. (2021) reported significant increases in both the generation (after 24hr) and survival (after 4 weeks) of new adult-born cells in 8-week-old mice in response to 3 months of ADF or 10% CR. In addition, BrdU^+^ cell number was not altered in GHS-R^−/−^ mice relative to WT mice (P = 0.1709), suggesting that GHS-R signalling does not enhance new cell generation in the DG. Notably, the BrdU^+^ cell number was significantly reduced in the adult mice in comparison to adolescent mice, suggesting an age-related decline in adult-born cell production (main effect of age, P < 0.0001). This finding is consistent with previous work that demonstrated a similar age-related decline in hippocampal cell division (Khun et al.1996).

**Figure 4.**
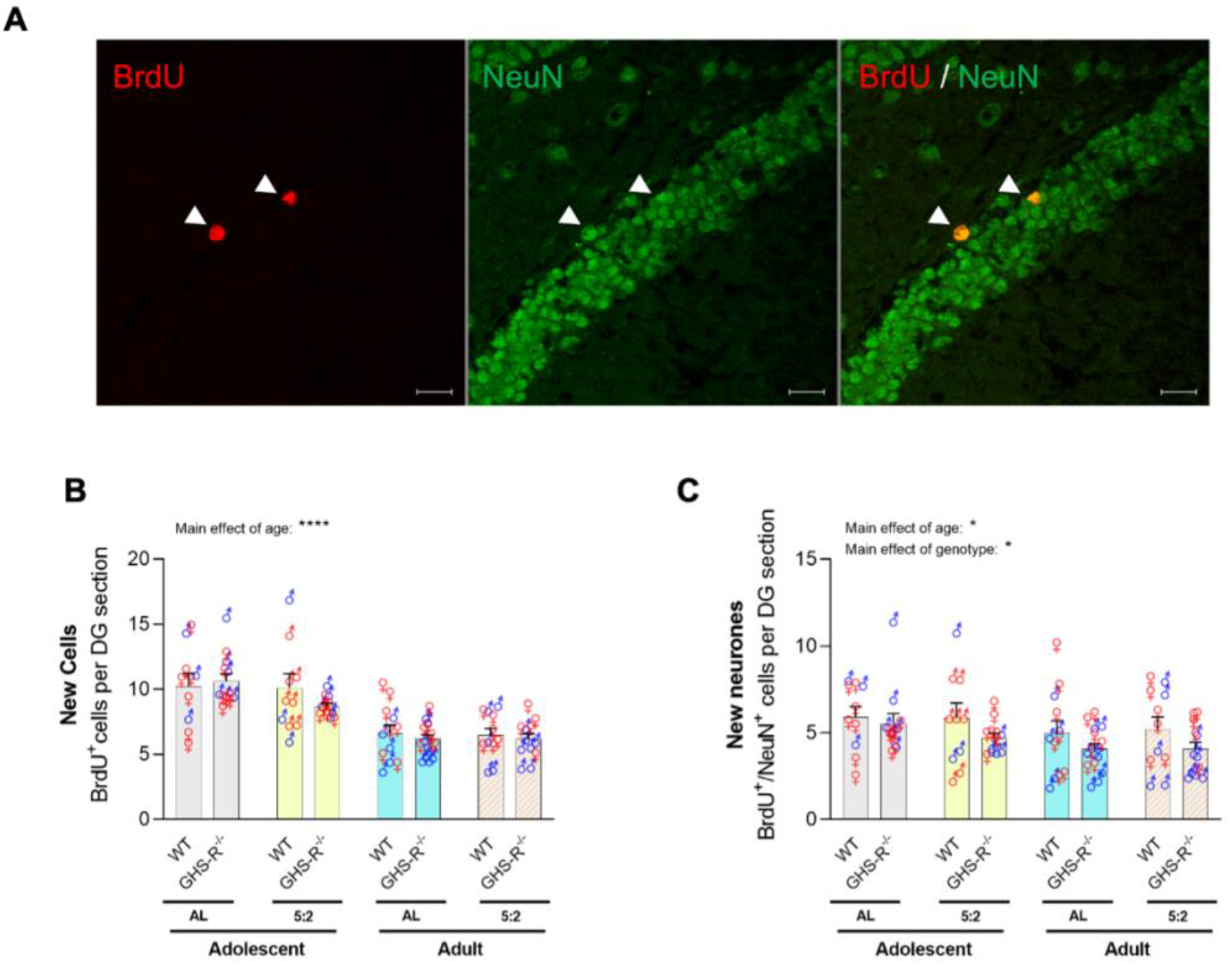
New adult born cells and neurones. Representative images of (A) new cells (BrdU) and new neurones (BrdU/NeuN) (scale bar = 20um). Quantification of new cells (B) and new neurones (C) in the adult hippocampus of adolescent and adult mice.

### The 5:2 diet did not alter the number of new adult-born neurones (BrdU^+^/NeuN^+^) in the hippocampal DG, but GHS-R was needed for maintaining neurogenesis

To determine the fate of the new BrdU^+^ cells, we explored the effect of the 5:2 diet compared to *ad-libitum* feeding on the generation of adult-born hippocampal neurones (BrdU^+^/NeuN^+^), in both adolescent and adult WT and GHS-R^−/−^ mice (Fig 4 A,C). BrdU^+^/NeuN^+^ immunoreactive cells in the SGZ and GCL were counted across the entire rostro-caudal extent of the hippocampus to reveal no main effect of diet on the number of new adult-born neurones (P = 0.9741). Notably, GHS-R^−/−^ mice had significantly fewer BrdU^+^/NeuN^+^ cells compared with WT mice across the adolescent and aged populations (main effect of genotype, P = 0.0082), suggesting a role for GHS-R1a signalling in the enhancement of AHN (Fig 4C). Moreover, BrdU^+^/NeuN^+^ cell numbers were significantly reduced in aged mice in comparison to their younger counterparts (main effect of age, P = 0.0359), supporting the notion of an age-related decline in AHN – as described previously by others (Khun et al.1996). There were no significant factor interactions, and post-hoc analyses revealed no statistically significant changes between groups. These findings were somewhat unexpected, as previous studies reported increased AHN in response to CR regimens. Indeed, our previous work demonstrated that a 2-week period of 30% CR increased the number of new adult-born neurones and the rate of neuronal differentiation in the hippocampal DG, and improved remote contextual fear memory in 12-week-old mice (Hornsby et al. 2016).

### The age-related decline in AHN was more pronounced in the suprapyramidal blade of the rostral DG

Given the distinct roles of new adult-born neurones in the rostral and caudal poles of the DG (Anacker et al., 2018), BrdU^+^/NeuN^+^ cells were quantified according to their location along the hippocampal rostro-caudal axis (also known as the dorso-ventral or septo-temporal axis). No significant differences were observed in the 5:2 *vs ad-libitum* fed mice in either the rostral (Table S2, P = 0.4773) or caudal (Table S2, P = 0.7724) DG sections. As before, a main effect of genotype was observed, with a significantly lower BrdU^+^/NeuN^+^ cell number in GHS-R^−/−^ mice compared with WT mice in both rostral (P = 0.0357) and caudal (P = 0.0195) DG sections. This suggests that neurogenic GHS-R signalling occurs at both poles of the DG. Interestingly, the adult mice exhibited a significant reduction in the number of new adult-born neurones compared to the adolescent mice in rostral (P = 0.0113), but not caudal (P = 0.0947) DG sections. Whilst the mean number of new adult-born neurones in the caudal DG sections was consistently decreased in the aged mice, this was not statistically significant, suggesting that the observed age-related decline in AHN is more pronounced in the rostral DG. No significant factor interaction effects were detected, and post-hoc analyses revealed no significant differences between groups. These data do not support the hypothesis that 5:2 diet stimulates AHN.

To provide further insight into the spatial organisation of new adult-born neurones, BrdU^+^/NeuN^+^ cells were quantified in the suprapyramidal and infrapyramidal blades of the DG in both rostral (Table S3-S4) and caudal (Table S5-S6) hippocampus. Dietary regimen had no significant effect on the BrdU^+^/NeuN^+^ cell number in the suprapyramidal (P = 0.8322) or infrapyramidal (P = 0.4221) blade in rostral DG sections. This was also the case for the suprapyramidal (P = 0.6524) and infrapyramidal (P = 0.4766) blades in caudal DG sections. Interestingly, the main effect of genotype on BrdU^+^/NeuN^+^ cell number remained significant in the suprapyramidal but not infrapyramidal blade of rostral DG sections (P = 0.0194 and P = 0.1237, respectively), and the infrapyramidal blade (P = 0.0209) but not suprapyramidal blade of caudal DG sections (P = 0.0616). New adult-born neurone numbers were significantly lower in adult mice compared to their adolescent counterparts in the suprapyramidal and infrapyramidal blades of rostral DG sections (P = 0.0297 and P = 0.0409, respectively) but only the suprapyramidal blade of the caudal DG (P = 0.0488). Notably, the rostral pole of the DG is particularly sensitive to both acyl-ghrelin and CR mediated increases in AHN (Kent et al.2015; Hornsby et al.2016). Combined, these findings identify the rostral DG as being particularly relevant to the age-related impairments in neurogenesis and cognition, and imply that ghrelin signalling may slow or even reverse age-related cognitive decline by activating neurogenic signalling in this region.

### The 5:2 diet did not increase the number of new neural stem cells (NSCs) in the hippocampal DG, but GHS-R was needed for regulating new NSC number

Next, to determine whether the 5:2 diet regulates NSC number in the hippocampal DG, BrdU^+^ immunoreactive cells in the SGZ were assessed for co-expression with Sox2 - a marker of type II NSCs. To allow new NSCs to be distinguished from a subset of new astrocytes that also express Sox2, immunoreactive cells were assessed for co-expression with the astrocytic marker, S100ß (Fig 5A). New NSCs (BrdU^+^/Sox2^+^/S100ß^−^) were counted across the entire rostro-caudal hippocampal SGZ (Fig 5B). Irrespective of age and genotype, there was no significant difference in the 5:2 fed mice compared with the *ad-libitum-*fed mice (P = 0.2599), suggesting that the 5:2 dietary regimen does not impact NSC number.

**Figure 5.**
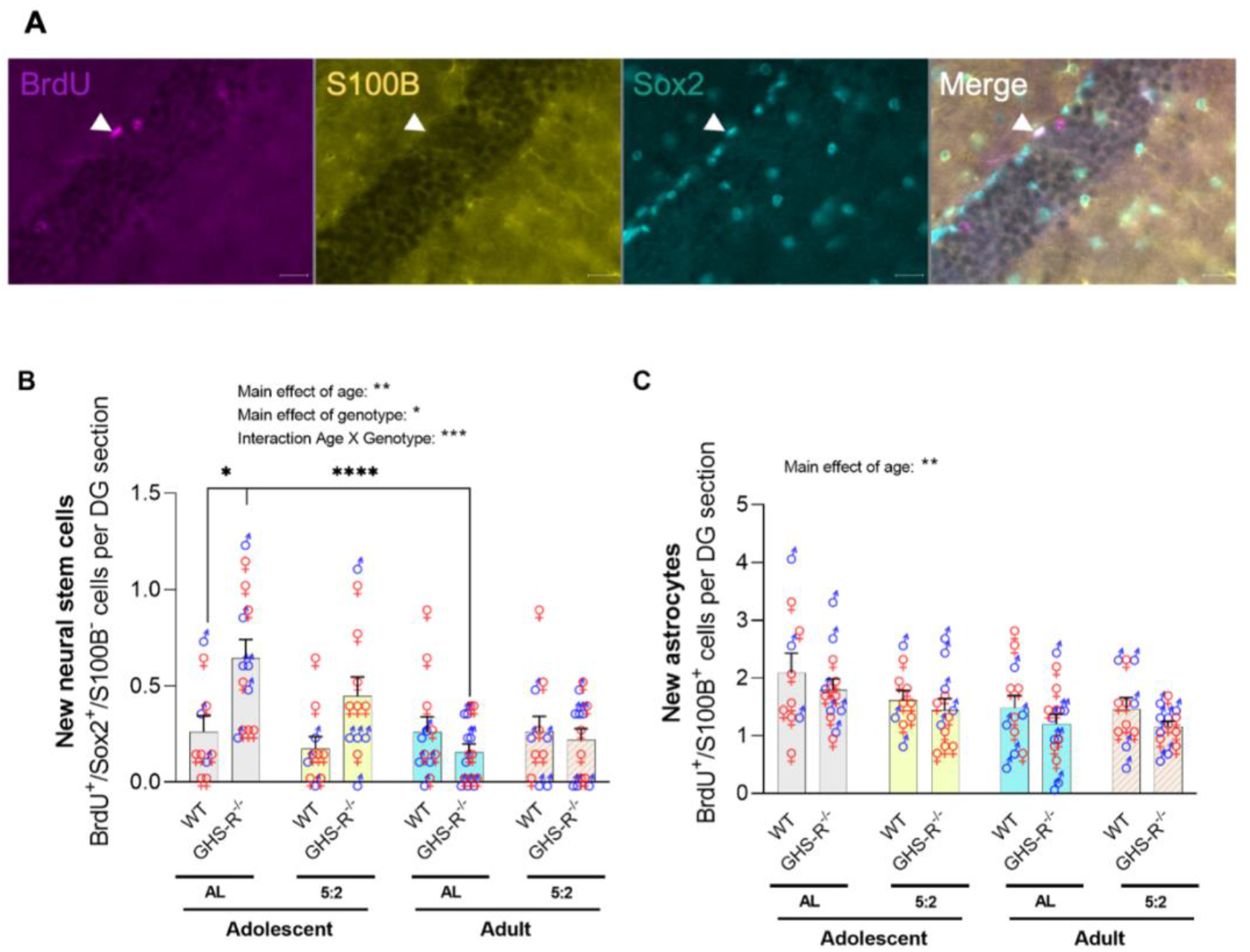
New neural stem cell and astrocytes. (A) Representative images of new neural stem cell (BrdU+/Sox2+/S100B−) and new astrocytes (BrdU+/S1008+) in adolescent and adult hippocampus (scale bar = 20um). (B) Graphs of new neural stem cell number (B) and new astrocytes (C).

Adult mice had fewer new NSCs (BrdU^+^/Sox2^+^/S100ß^−^) compared with their adolescent counterparts (Fig 4B; main effect of age, P = 0.0037), with post hoc analysis revealing a significant difference between the young and aged *ad-libitum* fed GHS-R^−/−^ mice (P = 0.0001), suggesting an age-related decline in new NSC number. While a main effect of genotype was observed (P = 0.0277), more notably, there was a highly significant interaction between genotype × age (P = 0.0003). In adolescent groups, GHS-R^−/−^ mice exhibited an increased number of new BrdU^+^/Sox2^+^/S100ß^−^NSCs in comparison to WT mice, with post-hoc analysis revealing a significant difference between the *ad-libitum* fed WT and GHS-R^−/−^ mice (P = 0.0184). Collectively, these findings suggest that the removal of GHS-R has a detrimental effect on the regulation of new NSC number in the neurogenic niche of the DG.

### New adult born NSCs were not differentially regulated by 5:2 diet in the rostral and caudal DG

To determine whether NSC renewal was differentially affected in the rostral and caudal poles of the DG, new BrdU^+^/Sox2^+^/S100ß^−^ NSCs were quantified across the rostro-caudal axis of the hippocampus. Consistent with the findings for the entire rostro-caudal axis, there was no main effect of dietary regimen on new BrdU^+^/Sox2^+^/S100ß^−^ NSC number when rostral (P = 0.929, Table S12) and caudal (P = 0.1584, Table S12) DG sections were analysed separately. The main effect of age remained significant in rostral (P = 0.0075) but not caudal (P = 0.053) poles, with post-hoc analyses revealing significant differences between the adolescent and adult groups in the *ad-libitum* fed GHS-R^−/−^ mice (rostral, P = 0.0016; caudal P = 0.0104). Furthermore, a significant interaction was reported between age and genotype (rostral, P = 0.0047; caudal, P = 0.0038), with more notable differences in the new BrdU^+^/Sox2^+^/S100ß^−^ NSC number between WT and GHS-R^−/−^ mice in the adolescent group, irrespective of feeding regimen (Table S12).

Our findings demonstrate that the 5:2 diet had no significant effect on new NSC number in the hippocampal SGZ (P = 0.2599). A previous study reported an increased number of NSCs and neural progenitor cells in the hippocampal DG in female mice following long-term exposure to 40% CR, suggesting that CR reduces the age-related decline in neural stem and progenitor cell division (Park et al. 2013). Similarly, a stepwise CR approach transiently increased NSC proliferation in the neurogenic niche of the sub-ventricular zone in young, but not aged mice (Apple et al., 2019). These results suggest that distinct DR regimens differentially regulate NSC homeostasis.

### The 5:2 diet did not increase the number of new adult-born astrocytes in the hippocampal DG

In addition to new NSCs, we also characterised the generation of new astrocytes (BrdU^+^/S100ß^+^) in the hippocampal DG (Fig 5C). As with the other new-born cell types, adult animals had significantly fewer new astrocytes compared to adolescent animals (P = 0.0044). However, no significant differences were reported in the number of new astrocytes between AL and 5:2-fed mice (P = 0.1168), or between WT and GHS-R^−/−^ mice (P = 0.0648). Thus, while the generation of new astrocytes in the hippocampal DG appears to be susceptible to age related decline, it does not appear to be affected by 5:2 diet or loss of GHS-R expression.

### Spatial memory performance was not enhanced by 5:2 diet, but GHS-R was needed for intact memory function

Finally, to assess whether the 5:2 diet affects hippocampal dependent spatial memory performance, we utilised the object in place (OIP) task (Fig 6A), as previously described (Evans et al.2020). Consistent with our BrdU/NeuN data, the 5:2 diet did not improve spatial memory performance (as determined by OIP discrimination index) in either adolescent or adult animals (main effect of diet: P = 0.4301; age × diet: P = 0.2359), while loss of GHS-R expression impaired spatial memory performance independent of both age and diet (main effect of genotype: P = 0.0050). However, in contrast to the BrdU/NeuN data, as well as the data for the other cellular markers of AHN (Ki67, DCX), a significant age-related decline in spatial memory performance was not detected with the OIP task (main effect of age: P = 0.3317) (Fig 6B).

**Figure 6.**
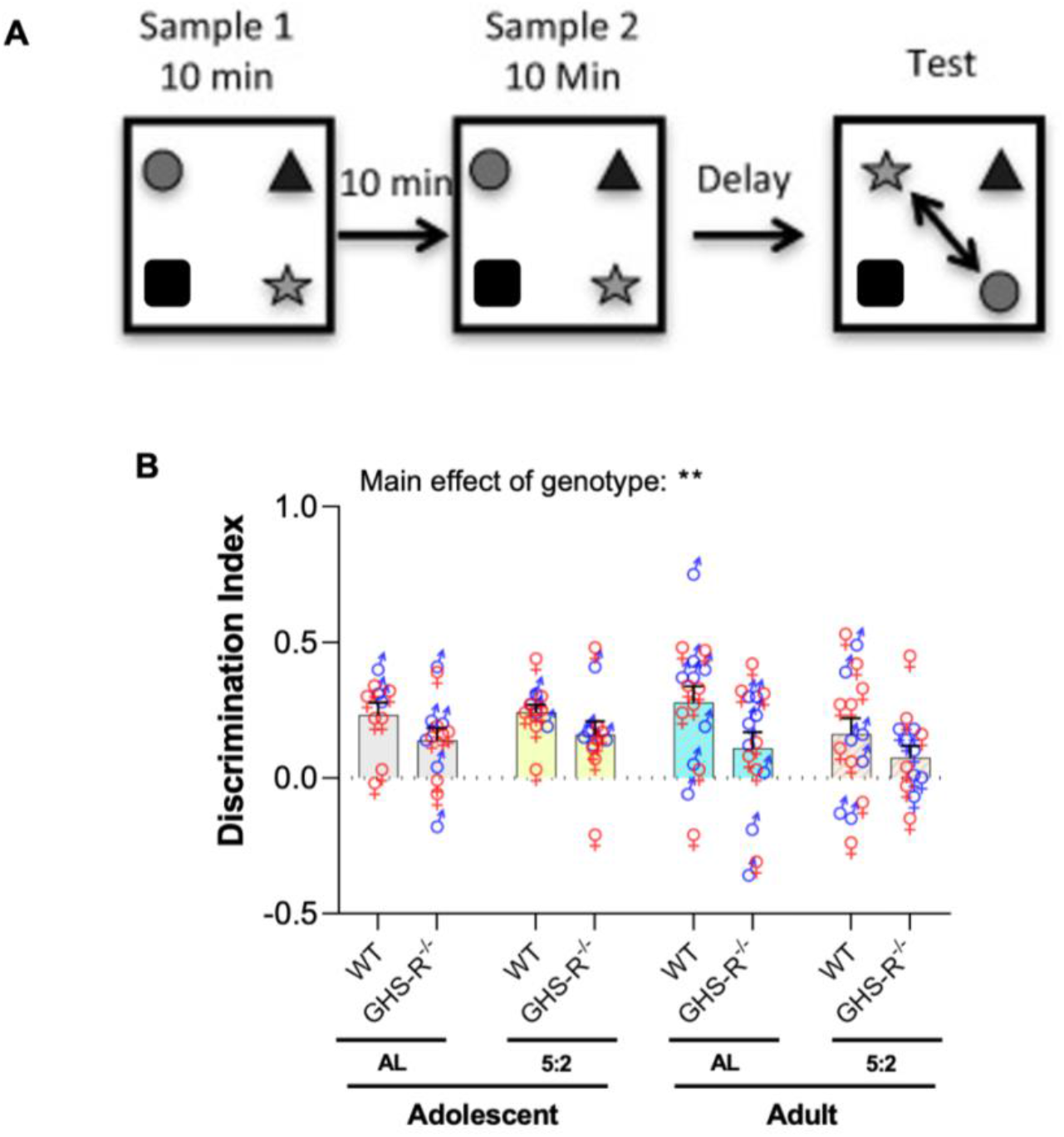
Spatial memory performance. (A) Schematic representation of the Object In Place (OIP) task to assess pattern separation performance. (B) Graph of the Discrimination Index of adolescent and adult mice in the OIP task.

## Discussion

Our findings relating to new adult-born hippocampal cell formation, differentiation and maturation in response to DR, which are distinct from other studies, may be attributable to differences in the regimens used. Previous rodent studies did not use a 5:2 feeding pattern and thus are not directly comparable to the present study. Those studies used more frequent DR periods, with rodents subjected to daily CR (Apple et al., 2019; Hornsby et al., 2016; Park et al. 2013) or variable IF regimens with a maximum of 24h between fasts (Baik et al., 2019; Cao et al., 2022; Dias et al., 2021; Kim et al., 2015; Lee et al., 2000, 2002). Indeed, ADF was implicated as a pro-neurogenic intervention two decades ago, whereby three months of ADF increased the survival of BrdU^+^ cells in the adult rodent DG three to four weeks after BrdU administration (Lee et al., 2000; Lee et al., 2002). Subsequently, several studies have reported that ADF (usually as a three-month intervention) stimulates aspects of neurogenesis (Baik et al., 2019; Cao et al., 2022; Dias et al., 2021; Kitamura et al., 2005 Kumar et al., 2009)). However, several of these studies limit their analysis to the proliferative stage (NSPCs and neuroblasts) without assessment of neuronal maturation, survival and functional integration into circuits regulating spatial memory (Cao et al., 2022; Kumar et al., 2009). Likewise, several studies report data for total BrdU^+^ cells in the DG, but not BrdU^+^/NeuN^+^ cells that are indicative of adult born neurones (Kitamura et al., 2005; Lee et al., 2000; Lee et al., 2002a; Lee et al., 2002b).

Nevertheless, the recent study of Dias et al., (2021), which utilised comprehensive IHC characterisations to evaluate the effects of three-months ADF or 10% daily CR on AHN, supported the earlier conclusion that IF, as well as CR, stimulates AHN. Indeed, both dietary regimens significantly increased total BrdU^+^ cells (24-hr and 4-weeks post injection), DCX^+^ cells, and the percentage of BrdU^+^/NeuN^+^ cells, compared to AL controls. Moreover, the number of BrdU^+^ cells at the 24-hr time point, as well as the number of DCX^+^ cells and the percentage of BrdU^+^/NeuN^+^ cells, was also significantly higher for the ADF group compared to the CR group; leading the authors to conclude that IF is more effective at stimulating AHN than daily 10% CR.

Our findings, utilising the 5:2 diet, do not support the notion that IF supports AHN. Despite reporting increased immature DCX^+^ neurone number, the 5:2 diet does not stimulate cell division, new mature neurone formation or functional integration of adult born neurones to support hippocampal dependent spatial memory. This contrasts with our previous study demonstrating that 2-weeks of daily 30% CR, followed by 2-weeks of *ad libitum* feeding, increased AHN (Hornsby et al., 2016). However, compared to ADF and continuous mild CR, the 5:2-fed mice in the present study underwent less frequent DR; being subjected to 2 non-consecutive fasting days every week (Mondays and Thursdays), with 2 to 3 days of *ad libitum* feeding between each fasting period. Thus, it is possible that the less frequent restrictive periods, with lengthened *ad libitum* feeding intervals in between, were insufficient to achieve the pro-neurogenic effects reported elsewhere. Another possible explanation for the distinct outcomes is the duration of the DR regimes. In the present study, mice were subjected to six weeks of the 5:2 diet, whereas many of the studies reporting improvements in aspects of neurogenesis exposed animals to 3 months or longer of IF, most commonly using an ADF method. Our previously published study is an exception, where mice were subjected to only 2 weeks of DR (Hornsby et al. 2016). However, this was in the form of 30% daily CR, which may be more efficient at enhancing AHN in the short-term due to the regime of daily restriction. Therefore, it is possible that this six-week period was an insufficient length of time for the 5:2 dietary regimen to elicit pro-neurogenic effects. Despite this, a study into the effect of IF on AHN-associated functions demonstrated that 4 weeks of the 5:2 diet was sufficient to increase pattern separation performance in a human population, whilst also decreasing memory recognition (Kim et al. 2020). These cognitive tests are an indirect measure of AHN and therefore further analysis is required to investigate the impact of the 5:2 diet on AHN at both the cellular and functional level.

Interestingly, a recent pre-print from Gabarró-Solanas et al., (2022) questions the suitability of ADF as a pro-neurogenic intervention, with neither 1-month or 3-months of ADF stimulating AHN. Utilising a tamoxifen inducible transgenic model (Glast-CreER^T2^;RYFP mice) to trace NSC lineage, ADF did not enhance NSC proliferation (as determined by Ki67 expression and Edu incorporation), nor the abundance of immature neurones (DCX^+^ cells). Moreover 3-months of ADF failed to increase the number of new adult-born neurones (NeuN^+^/YFP^+^), while 1 month of ADF significantly reduced the number of new adult-born neurones compared to AL control. Given that most studies on AHN utilise the C57BL/6J strain of mice, the use of the GlastCreER^T2^;RYFP model may have underscored these observations. Also, tamoxifen has been reported to impair various physiological processes, including neurogenesis (Smith et al., 2022). However, to account for these potentially confounding factors, they characterised the effect of their ADF regimen in C57BL/6 mice, as well as potential anti-neurogenic effects arising from tamoxifen administration to report no data supporting ADF as a pro-neurogenic intervention.

Collectively, our findings and those of Gabarró-Solanas et al., (2022) do not align with other studies on DR and neurogenesis. However, variations in the age and sex of animals used in each of these studies limit the value of direct comparisons. Thus, further comprehensive studies are required to determine the optimal level and duration of DR restriction, as well as optimal frequency for restrictive periods, for the enhancement of AHN.

We observed a significant reduction in the number of new adult-born cells (BrdU^+^) in the DG of aged mice compared with young mice (P < 0.0001). This is in accordance with many previous studies showing that the detrimental effect of ageing on the generation of new adult-born cells in the hippocampal DG (Kuhn et al., 1996; Bondolfi et al., 2004). Immature neurone (DCX^+^) and new adult-born neurone (BrdU^+^/NeuN^+^) production was also significantly decreased in the aged mice in comparison to the young mice (P < 0.0001; P = 0.0359, respectively), supporting the notion that age is a key modulator of AHN (Kempermann et al., 1998; Kuhn et al., 2018). The present findings also showed that this age-related decline was more pronounced in the rostral DG. Indeed, the rostral pole of the DG is implicated in cognitive functions such as learning, memory and spatial navigation (Faneslow and Dong, 2010). Moreover, the suprapyramidal blade is thought to contain a greater proportion of active granule cells, with an increased responsiveness during spatial tasks (Gallitano et al., 2016; Schmidt et al., 2012). This raises the question of whether the reduction in new adult-born neurones in the rostral DG, particularly in the suprapyramidal blade, leads to age-related cognitive decline and spatial memory defects (Bach et al., 1999; Ngwenya et al., 2015).

The age-associated decrease in new neurone production may be attributable to alterations in NSC homeostasis (Ibrayeva et al., 2021). Our findings demonstrate an age-related decline in new NSC number in the SGZ across the entire rostro-caudal axis (P = 0.0037). This is consistent with a significant decline in the number of new NSCs in the SGZ of the aged rat brain (Hattiangadya & Shettya 2008). Moreover, a decrease in the number of Sox2^+^ quiescent neural progenitors (QNPs) in the DG of aged humans was reported as being specific to the anterior-mid DG (rostral) rather than across the entire rostro-caudal axis (Boldrini et al., 2018). These findings suggest that this age-related decline in NSC renewal is conserved across different mammalian species.

We also sought to determine the effect of GHS-R signalling on new NSC number. Our previous study reported that exogenous acyl-ghrelin, administered intraperitoneally at a physiological dose, did not enhance new hippocampal NSC number in 3-month-old female rats (Kent et al.2015). Here, we show that new NSC number was increased in adolescent GHS-R^−/−^ mice in comparison to their wild-type counterparts, with the loss of this effect in adult mice. This increase is unlikely to be a direct effect as GHS-R is not expressed in type II (Sox2^+^) NSCs in the SGZ (Hornsby et al., 2016). Instead, it may be due to an indirect mechanism yet to be elucidated, whereby the absence of GHS-R signalling promotes NSC renewal and/or possibly hinders NSC differentiation. Indeed, we have shown that the absence of GHS-R signalling reduces the rate of neuronal differentiation (Hornsby et al.2016). However, despite this increase in new NSCs during adolescence there was no corresponding increase in the number of new adult-born neurones at this age. These observations collectively suggest that the loss of GHS-R signalling leads to a shift in AHN dynamics, whereby the number of cells displaying early-maturation stage phenotypes is increased and mature neuronal phenotypes are decreased. Interestingly, the number of new adult-born NSCs was similar in adult WT and GHS-R^−/−^ mice, suggesting that the effect was limited to the developmental adolescent period.

The ‘disposable stem cell’ model suggests that activated NSCs undergo a series of rapid asymmetric divisions leading to neuronal differentiation and cell elimination coupled with the eventual differentiation of NSCs into astrocytes (Encinas et al. 2011). The in vivo activation of NSCs results in a burst of neurogenic activity, whereby self-renewal capacity is limited, and activated cells are subsequently lost from the NSC pool (Pilz et al.2018). These findings may provide an explanation for the age-related decline in NSC renewal. In contrast, other studies describe the long-term maintenance of hippocampal NSCs, whereby NSCs slowly alternate between periods of quiescence and activity, have prolonged self-renewal capabilities, and can generate new neurones and astrocytes (Bonaguidi et al. 2011; Dranovsky et al., 2017). Heterogeneous NSC populations have been identified with short-term and long-term self-renewal capabilities, each giving rise to new hippocampal neurones, even in the aged brain (Bottes et al., 2021; Ibrayeva et al., 2021). Notably, longer-lived NSC populations exhibit increased quiescence with successive cell divisions, leading to disturbed homeostasis and fewer NSCs in the aged brain (Harris et al., 2021; Ibrayeva et al., 2021; Kalamakis et al, 2019). Thus, the age-related decline in NSC renewal could instead be due to the altered quiescence of NSCs.

While the effect of GHS-R signalling on NSC renewal remains unclear, our findings, together with those from previous studies, suggest that GHS-R signalling enhances AHN by inducing the maturation of new adult-born neurones. Evidence from previous studies showed that AHN is decreased during ageing and dysregulated in neurodegenerative diseases (Kuhn et al., 1996; Moreno-Jimenez et al., 2019; Terreros-Roncal et al., 2021). Thus, future work into the mechanisms underlying the pro-neurogenic effects of GHS-R signalling may reveal new therapeutic targets for age-related decline and neurodegenerative diseases.

In summary, we confirm an age-related decline in AHN and that GHS-R is needed for intact AHN and spatial pattern separation performance. We show for the first time that GHS-R is essential for regulating new NSC formation in the hippocampal neurogenic niche. However, our data demonstrate that the 5:2 diet does not increase the formation of new adult born neurones or enhance hippocampal spatial memory performance. These findings suggest that distinct DR regimens differentially regulate neurogenesis in the adult hippocampus and that further studies are required to identify optimal protocols to support cognition during ageing.

## Methods

All animal work was performed at Cardiff University under the appropriate Home Office Animal Act 1986 approval.

### Mice

Homozygous loxTB-GHSR mice (herein referred to as GHS-R^−/−^) and age/sex-matched wild-type (WT) mice from the same colony (a gift from Professor Jeffrey Zigman, UT Southwestern; (Zigman et al., 2005)) were bred from mice heterozygous for the recombinant lox-P flanked transcriptional blockade of the *Ghsr* allele. Mice belonged to one of two age groups: adolescent (7-weeks old at the start of the study; N=45), or adult (7-month-old; N=51). Mice were housed at Cardiff University under standard laboratory conditions on 12h light: 12h dark cycles (lights on at 06:00 h). For the labelling of proliferating cells, all mice were given twice daily intraperitoneal injection of the thymidine analogue, BrdU (50mg/kg) on days 1 and 2 of the study.

Young and aged mice populations were each divided into four groups (giving 8 groups in total, n=10-15 per group) based on genotype and dietary regimen: *ad libitum* fed WT, 5:2-fed WT, *ad libitum*-fed GHSR^−/−^ and 5:2-fed GHSR^−/−^. Mice were fed standard laboratory chow (Harlan Laboratories) and maintained on either *ad libitum* (unrestricted) or 5:2 dietary regimens for six weeks. The 5:2 regimen consisted of two non-consecutive days without food access (Monday and Thursday), and *ad libitum* feeding on the remaining five days, with access to food provided at 4pm, 2h prior to lights off. Body weight was measured between 9-10am on Mondays, Wednesdays, and Fridays. After this six-week period, all mice were subjected to one week of *ad libitum* feeding whilst behavioural tests were conducted.

Mice were sacrificed by intracardial perfusion with 4% paraformaldehyde (PFA). Brains were removed and post-fixed in 4% PFA overnight at 4°C, prior to cryoprotection in 30% sucrose solution. Brains were transported to Swansea University and stored at −80°C. Brains were cut into 30μM-thick coronal sections along the rostro-caudal extent of the hippocampus using a freezing-stage Microtome (MicroM HM450, ThermoScientific). Sections were collected 1:12 in 0.1% sodium azide in PBS and stored at 4°C until required for immunohistochemistry.

### Immunohistochemistry

For immunofluorescent analysis of BrdU^+^/Dcx^+^, sections were washed three times in PBS for 5 minutes, permeabilized in methanol at −20°C for 2 minutes and washed (as before) prior to pre-treatment with 2N HCl for 30 minutes at 37°C followed by washing in 0.1 M borate buffer, pH8.5, for 10 minutes. Sections were washed as before and blocked with 5% normal donkey serum (NDS) in PBS plus 0.1% triton (PBS-T) for 60 minutes at room temperature. Sections were incubated overnight at 4°C in rat anti-BrdU (1:400, AbD Serotec) and goat anti-Dcx (1:200, Santa Cruz Biotechnology, USA) diluted in PBS-T. Tissue were washed as before and incubated in donkey anti-rat AF-488 (1:500, Life Technologies, USA) and donkey anti-goat AF-568 (1:500, Life Technologies, USA) in PBS-T for 30 minutes in the dark. After another wash sections were mounted onto superfrost+ slides (VWR, France) with prolong-gold anti-fade solution (Life Technologies, USA).

For immunofluorescent analysis of Sox2, sections were treated as above with the exception of antigen retrieval being performed in sodium citrate at 70°C for 1h (rather than 2N HCl or borate buffer) with subsequent blocking in 5% normal goat serum (NGS). Immunoreactivity was detected using rabbit anti-Sox2 (1:500, ab97959, Abcam) and goat anti-rabbit AF-568 (Life Technologies, USA). Nuclei were counterstained with Hoechst prior to mounting as described above.

For immunofluorescent analysis of Sox2^+^/ S100β^+^, sections were washed three times in PBS for 5 minutes, permeabilised in methanol for 2 minutes at −20°C and washed again (as above). Antigen retrieval was performed with sodium citrate buffer for 1 hour at 70°C, sections were washed as before and subsequently blocked with 5% normal goat serum (NGS) in PBS+0.1% Triton-X (PBS-T) for 1 hour at room temperature. Sections were incubated overnight at 4°C in rabbit anti-Sox2 (1:1000, ab97959, Abcam) diluted in PBST. Sections were washed as before and incubated in goat anti-rabbit AF-568 (1:500, Life Technologies) in PBST for 30 minutes at room temperature, in the dark. Following another wash step, sections were incubated for 1 hour at room temperature with mouse anti-S100β (1:1000, S2532, Sigma) in PBST. Following another set of washes, sections were incubated in goat anti-mouse AF-488 (1:500, Life Technologies) in PBST for 30 minutes at room temperature and protected from light. After a final wash, sections were mounted onto Superfrost^+^ Plus slides (VWR) with Prolong-gold anti-fade mounting solution (Invitrogen) and coverslipped.

For DAB-immunohistochemical analysis of Ki67 and DCX labeling, sections were washed in 0.1M PBS (2×10mins) and 0.1M PBS-T (1×10 mins). Subsequently, endogenous peroxidases were quenched by washing in a PBS plus 1.5% H_2_O_2_ solution for 20 minutes. Sections were washed again (as above) and incubated in 5% NGS in PBS-T for 1h. Sections were incubated overnight at 4°C with rabbit anti-Ki67 (1:500, ab16667, Abcam) or guinea pig anti-DCX (1:15,000 Sigma AB2253), in PBS-T and 2% NGS solution. Another wash step followed prior to incubation with biotinylated goat anti-rabbit (1:500 Vectorlabs, USA) for Ki67 or biotinylated goat anti-guinea pig (1:500 Vectorlabs, USA) for DCX, in PBS-T for 70 minutes. The sections were washed and incubated in ABC (Vectorlabs, USA) solution for 90 minutes in the dark prior to another two washes in PBS, and incubation with 0.1M sodium acetate pH6 for 10 minutes. Immunoreactivity was developed in nickel-enhanced DAB solution followed by two washes in PBS. Sections were mounted onto superfrost^+^ slides (VWR, France) and allowed to dry overnight before being de-hydrated and de-lipified in increasing concentrations of ethanol. Finally, sections were incubated in Histoclear (2×3 mins; National Diagnostics, USA) and coverslipped using Entellan mounting medium (Merck, USA). Slides were allowed to dry overnight prior to imaging.

### Immunofluorescence

All immunofluorescence experiments were carried out on free-floating sections at room temperature, unless stated otherwise.

### Double Sequential Immunofluorescence (BrdU/NeuN)

Four rostral and four caudal brain sections were used for each mouse. Sections were washed three times in 1X Phosphate Buffered Saline (PBS) on a rocker for 5 min, followed by permeabilisation in methanol at −20°C for 2 min, and subsequent washing in PBS as before. For antigen retrieval, sections were treated with 2M HCl at 37°C for 30 min. This was followed by washing in 0.1M borate buffer (pH 8.5) for 20 min. Sections were washed in PBS as before, prior to blocking with 5% Normal Goat Serum (NGS) in PBS plus 0.1% Triton-X100 (PBS-T) for 1 h. Excess block was removed, and sections were incubated overnight at 4°C in Rat anti-BrdU (MCA2060, AbD Serotec) diluted 1:400 in PBS-T. Sections were washed in PBS as before, and incubated for 1 h in the dark in goat anti-rat AF488 (A-11006, Life Technologies, USA) diluted 1:500 in PBS-T. All washes and incubations following this step were carried out in the dark. Following another series of washes in PBS, sections were treated with mouse anti-NeuN (MAB377, Millipore, USA) diluted 1:1000 in PBS-T and incubated for 1 h. Three further washes in PBS were carried out, followed by incubation for 30 min in Goat anti-mouse AF568 (A-11004, Life Technologies, USA) diluted 1:500 in PBS-T. Sections were washed once in Hoescht nuclear stain diluted 1:5000 in PBS for 5 min, followed by two additional washes in PBS as before. Sections were mounted onto superfrost^+^ slides with Prolong Gold anti-fade solution (Life Technologies, USA) equilibrated to room temperature. Coverslips were applied and slides were left covered on the bench for 24 h to allow to dry, prior to storing at 4°C in the dark.

### Triple Immunofluorescence (BrdU/Sox2/S100b)

For BrdU/Sox2/S100b immunofluorescence, two rostral and two caudal brain sections were used from each mouse. Initially, sections were treated as previously described for BrdU/NeuN, with the exception that non-specific binding sites were first blocked in 5% Normal Donkey Serum (NDS) in PBS-T for 30 min, and then in 5% NGS for 30 min. In addition, the primary antibodies were applied together in a cocktail that consisted of rat anti-BrdU (1:400), rabbit anti-Sox2 (1:500; ab97959, Abcam) and mouse anti-S100b (1:500; S2532, Sigma) in PBS-T, and incubated overnight at 4°C. Sections were then washed in PBS as before. This was followed by incubation for 30 min in the dark with the secondary antibody cocktail that consisted of goat anti-rat AF488, donkey anti-rabbit AF647 (A-31573, Life Technologies, USA) and goat anti-mouse AF568, all diluted 1:500 in PBS-T. Sections were washed once in Hoescht nuclear stain in PBS and twice more in PBS, and then mounted as previously described in section 2.3.1.

### Quantification of immunoreactive cells

Immuno-stained brain tissue was imaged by light microscopy (Nikon 50i) (for DAB), fluorescent (Axioscope, Zeiss) or confocal microscopy (LSM710 META, Zeiss). DAB-immunolabelled cells were quantified using Image J software (Ki67) or manually (DCX). Resulting numbers were divided by the number of coronal sections analyzed and expressed as cells per DG section. All analyses were performed blind to feeding pattern, genotype and treatment.

Immuno-histologically stained sections from each mouse were imaged using a fluorescent microscope (Axioscope, Zeiss or CLSM 810, Zeiss). A ×40 objective lens was used to manually count BrdU^+^ immunoreactive cells through the *z*-axis across the rostro-caudal extent of the GCL and SGZ. The SGZ was defined as the area occupied by approximately two cell bodies, above and below, the inferior edge of the GCL. To gain a further insight into the spatial organisation of immunoreactive cells, their location in either the suprapyramidal (upper) or infrapyramidal (lower) blade of the DG was specified.

Manual cell counting techniques are prone to variations in human perception, therefore, to maintain consistency, quantification for the BrdU/NeuN element of this study was performed by one operator. For the quantification of new mature neurones, BrdU^+^ immunolabelled cells were assessed for co-expression with the mature neuronal marker, Neuronal Nuclei (NeuN).

For the quantification of new stem cells, BrdU^+^ immunolabelled cells were assessed for co-expression with Sox2 - a marker of type II NSCs. Since Sox2 is also expressed by a subset of hippocampal astrocytes (Komitova & Eriksson, 2004), the astrocytic marker, S100b, was used to differentiate new stem cells (BrdU^+^/Sox2^+^/S100b^−^) from new astrocytes (BrdU^+^/Sox2^+^/S100b^+^). Newly formed double-positive astrocytes (BrdU^+^/Sox2-/S100b^+^) were also quantified. Cells labelled solely with BrdU (BrdU^+^/Sox2^−^/S100b^−^) were classed as new ‘other cells’ since it was not possible to definitively confirm the identities of these cells.

Resulting numbers were divided by the number of DG sections analysed to determine the mean number of immunoreactive cells per DG section for each mouse. For BrdU/NeuN analyses, the relative proportion of BrdU^+^ cells differentiated into mature neurones (BrdU^+^/NeuN^+^) were also determined for each mouse. All experimental and quantification steps were performed blinded to the feeding group and genotype of the mice to prevent bias. Representative images were acquired using the ZEN blue software and processed using ZEN lite.

### Object In Place Task

The apparatus used for all experiments was a large Perspex arena, 60 × 60 × 40 cm, with a pale grey floor and clear walls, which for the purpose of this experiment were covered with white paper. The box was placed on a square table at waist height. The apparatus was set up in a quiet and brightly lit (38 cd/m^2^ at the arena surface) behavioural testing room.

Exploration was recorded with an overhead camera. The camera input was used to monitor activity in the arena on a television monitor and each session was recorded using a Philips DVDR recorder.

The duration of object exploration throughout the trials was recorded manually with a stopwatch. All objects used were everyday objects made of non-porous materials. All objects were at least 10 cm high to avoid the mice climbing and sitting on the objects, and were all weighted so that they could not be displaced by the animals. Both the arena and the objects were cleaned thoroughly with water and ethanol wipes in between each trial to prevent the use of odour cues, urine and excrement were also removed from the arena after each trial.

The main aim of this experiment was to assess whether experimental conditions influenced memory for specific object-location associations.

The week prior to testing, mice were handled for 5 minutes a day, three times a week. For three days prior to testing, mice were placed in the behavioural test room in their home cages, for 30 minutes a day. On each of these days mice were also given habituation session in which to freely explore the arena with no objects present for 10 minutes. Training commenced the following day.

Mice were placed in the centre of the arena and presented with four different objects, each in a different corner of the arena during the sample and test trials. Mice were allowed to explore the arena and the objects for 10 minutes before being removed for a 10-minute interval spent in their home cage (located in the testing room). Mice were then given a second 10-minute sample phase. Following the second sample phase, the mice were returned to their home cage for a 10-minute retention interval (located in the testing room). In all experiments, mice received a 10-minute test phase following this interval. In the test phase, two of the objects swapped their spatial locations. This resulted in two familiar objects located in different positions, and two familiar objects that remained in their original location. The objects that exchanged their spatial locations were counterbalanced, and the location of the objects in the arena was also counterbalanced to avoid spatial biases.

For each experiment, the dependent variable was the amount of time spent by the animals exploring objects. Object exploration was defined as the time spent attending to (actively sniffing or interacting with) the object at a distance no greater than 1 cm. Object exploration was not scored if the animal was in contact with but not facing the object, or if it attempted to climb on the objects to look around the rest of the arena. In order to ensure that procedures were sensitive to differences between the groups independent of variation in individual contact times, a discrimination ratio was calculated for each experimental test phase. Discrimination ratios were calculated as follows; DR = average time exploring the two objects in new locations / (average time exploring the two objects in the familiar location + average time exploring the two objects in different locations).

### Statistical analysis

Statistical analyses were performed using GraphPad Prism 9.3.1. Statistical significance was assessed using a 3-way Analysis of Variance (ANOVA), with Tukey’s multiple comparisons test. Residuals were assessed for deviation from a normal distribution using a Shapiro-Wilk test. For all statistical tests, *, p < 0.05; **, p < 0.01; ***, p < 0.001; ****, p < 0.0001 were considered significant.

## Acknowledgments

This project was supported by a grant from the British Society of Neuroendocrinology to LR, TW and JSD, a grant from The Waterloo Foundation and The Rosetrees Trust to AKEH, JSD and TW. AKEH and TW wish to acknowledge the generous financial support of Cardiff University.

---

**Supplementary Figure.S1.**
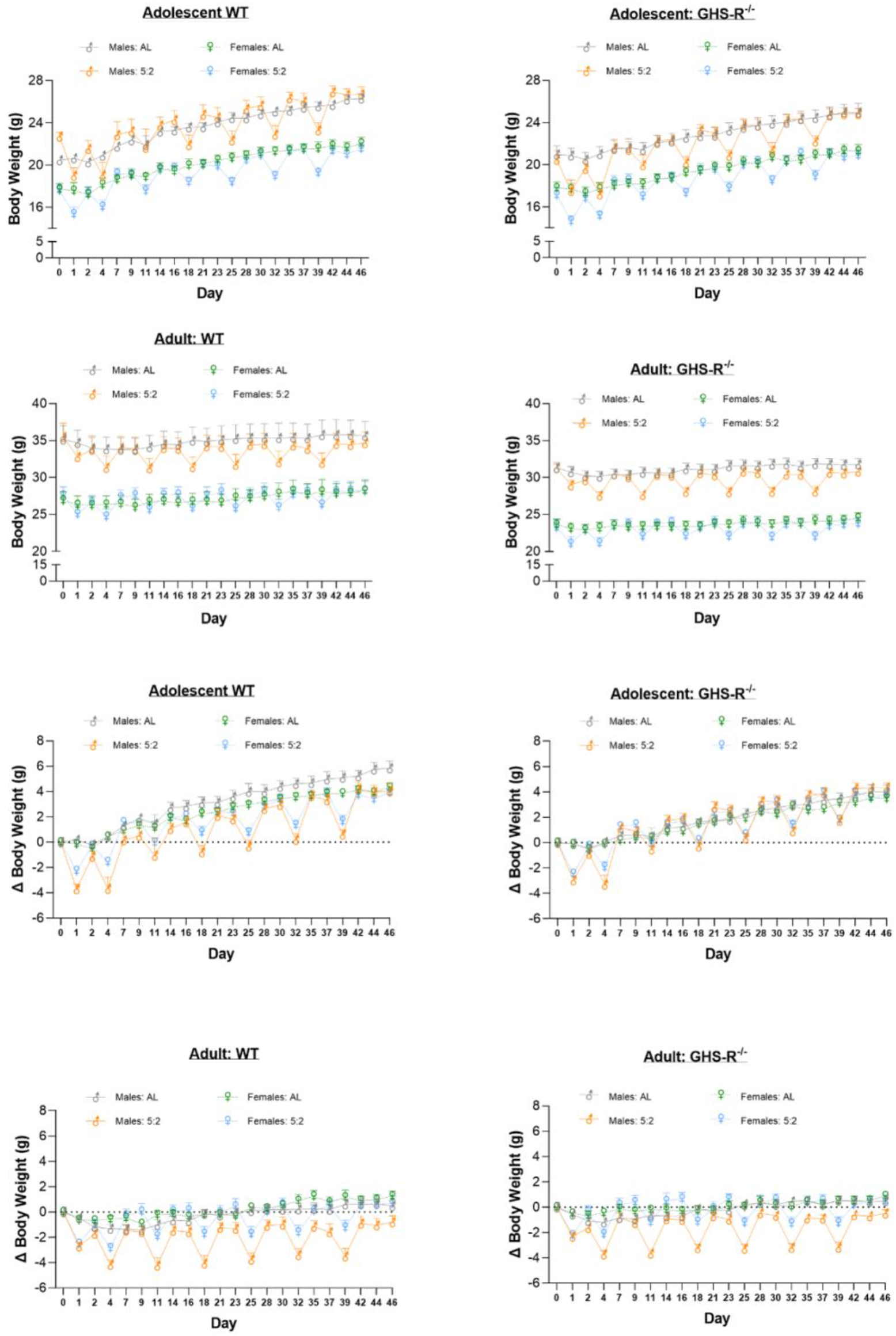
Males vs Females Repeated Measures.

## Supplementary Data

**Table S1.** New-born neurones (BrdU^+^/NeuN^+^) in the rostral and caudal poles of the hippocampal dentate gyrus (DG). Data expressed as mean counts (± SEM) per DG section.

|  | WT: AL | GHS-R<sup>-/-</sup> :<br>AL | WT: 5:2 | GHS-R<sup>-/-</sup> :<br>5:2 |
| --- | --- | --- | --- | --- |
| **Adolescent** |  |  |  |  |
| **Rostral** | 6.54 ± 0.83 | 5.30 ± 0.41 | 5.88 ± 1.11 | 4.89 ± 0.38 |
| **Caudal** | 5.21 ± 0.60 | 5.26 ± 0.51 | 5.85 ± 0.68 | 4.56 ± 0.27 |
| **Adult** |  |  |  |  |
| **Rostral** | 4.96 ± 0.82 | 4.00 ± 0.36 | 5.15 ± 0.66 | 4.08 ± 0.33 |
| **Caudal** | 4.96 ± 0.61 | 3.95 ± 0.36 | 5.39 ± 0.75 | 4.02 ± 0.49 |

**Table S2.**
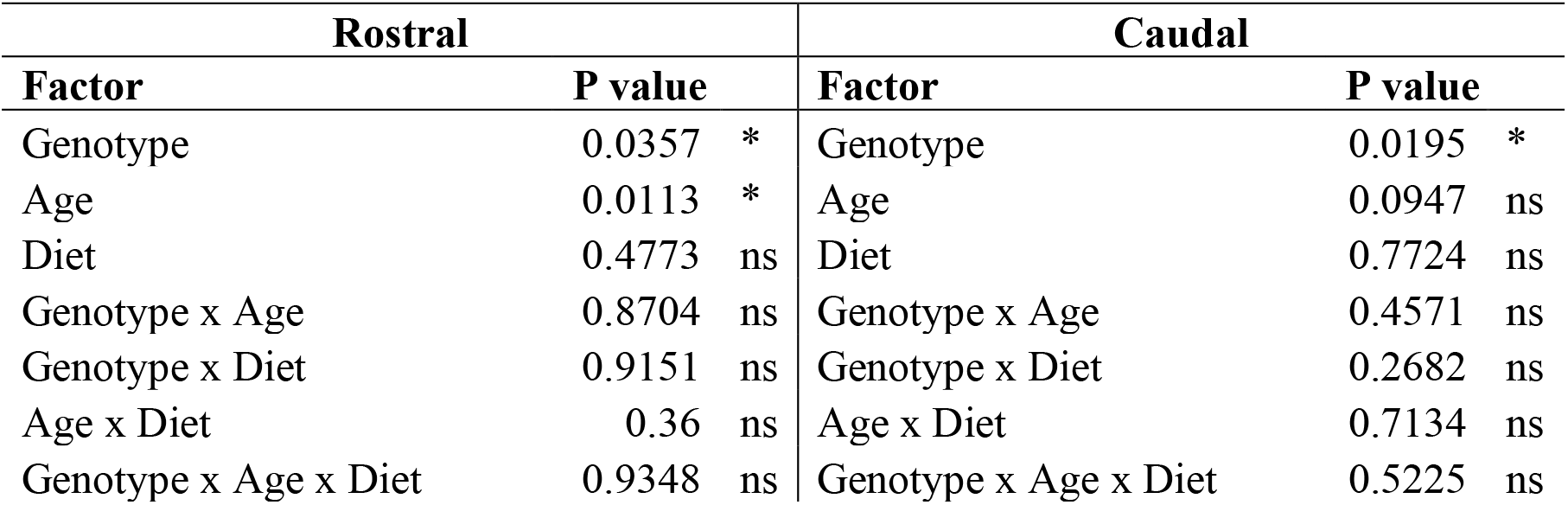
Summary of three-way ANOVA statistical analyses of new-born neurones (BrdU^+^/NeuN^+^) in the rostral and caudal poles of the hippocampal dentate gyrus (DG).

**Table S3.**
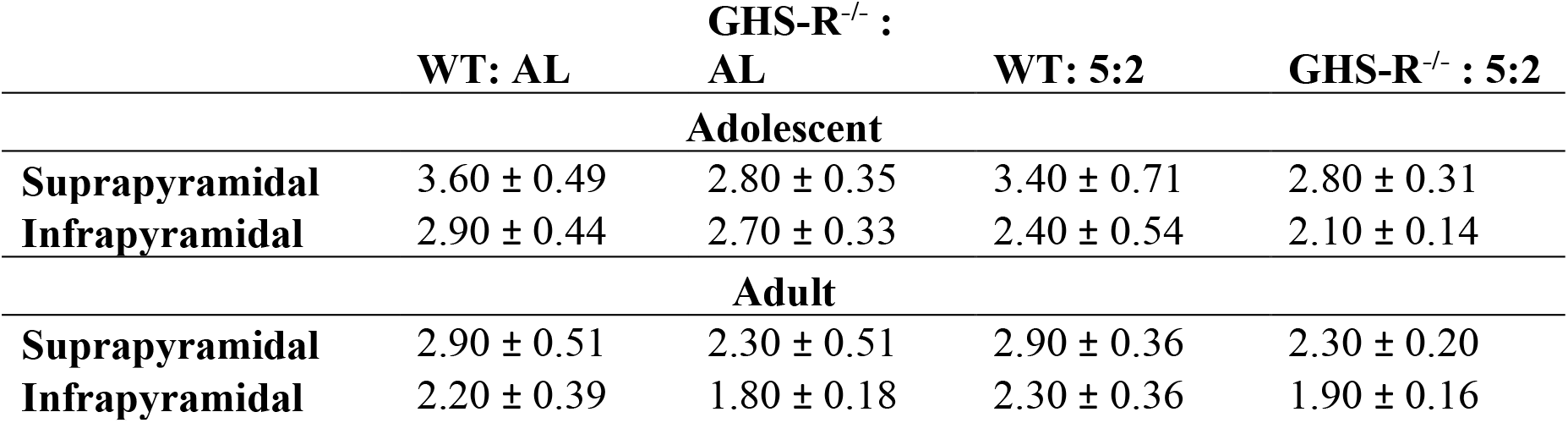
New-born neurones (BrdU^+^/NeuN^+^) in the suprapyramidal and infrapyramidal blades of the rostral dentate gyrus (DG). Data expressed as mean counts (± SEM) per DG section.

**Table S4.** Summary of three-way ANOVA statistical analyses of new-born neurones (BrdU^+^/NeuN^+^) in the suprapyramidal and infrapyramidal blades of the rostral dentate gyrus (DG).

| **Suprapyramidal** |  |  | **Infrapyramidal** |  |  |
| --- | --- | --- | --- | --- | --- |
| **Factor** | **P value** |  | **Factor** | **P value** |  |
| Genotype | 0.0194 | * | Genotype | 0.1237 | ns |
| Age | 0.0297 | * | Age | 0.0409 | * |
| Diet | 0.8322 | ns | Diet | 0.4221 | ns |
| Genotype x Age | 0.91 | ns | Genotype x Age | 0.8565 | ns |
| Genotype x Diet | 0.8367 | ns | Genotype x Diet | 0.8955 | ns |
| Age x Diet | 0.8974 | ns | Age x Diet | 0.1362 | ns |
| Genotype x Age x Diet | 0.9019 | ns | Genotype x Age x Diet | 0.9045 | ns |

**Table S5.** New-born neurones (BrdU^+^/NeuN^+^) in the suprapyramidal and infrapyramidal blades of the caudal dentate gyrus (DG). Data expressed as mean counts (± SEM) per DG section.

|  | GHS-R<sup>-/-</sup> : |  |  |  |
| --- | --- | --- | --- | --- |
|  | WT: AL | AL | WT: 5:2 | GHS-R<sup>-/-</sup> : 5:2 |
| Adolescent |  |  |  |  |
| Suprapyramidal | 3.10 ± 0.39 | 3.20 ± 0.42 | 3.30 ± 0.34 | 2.50 ± 0.21 |
| Infrapyramidal | 2.10 ± 0.26 | 2.20 ± 0.23 | 2.50 ± 0.38 | 1.90 ± 0.11 |
| Adult |  |  |  |  |
| Suprapyramidal | 2.80 ± 0.35 | 2.30 ± 0.21 | 2.90 ± 0.39 | 2.30 ± 0.28 |
| Infrapyramidal | 2.10 ± 0.28 | 1.70 ± 0.14 | 2.50 ± 0.36 | 1.60 ± 0.30 |

**Table S6.**
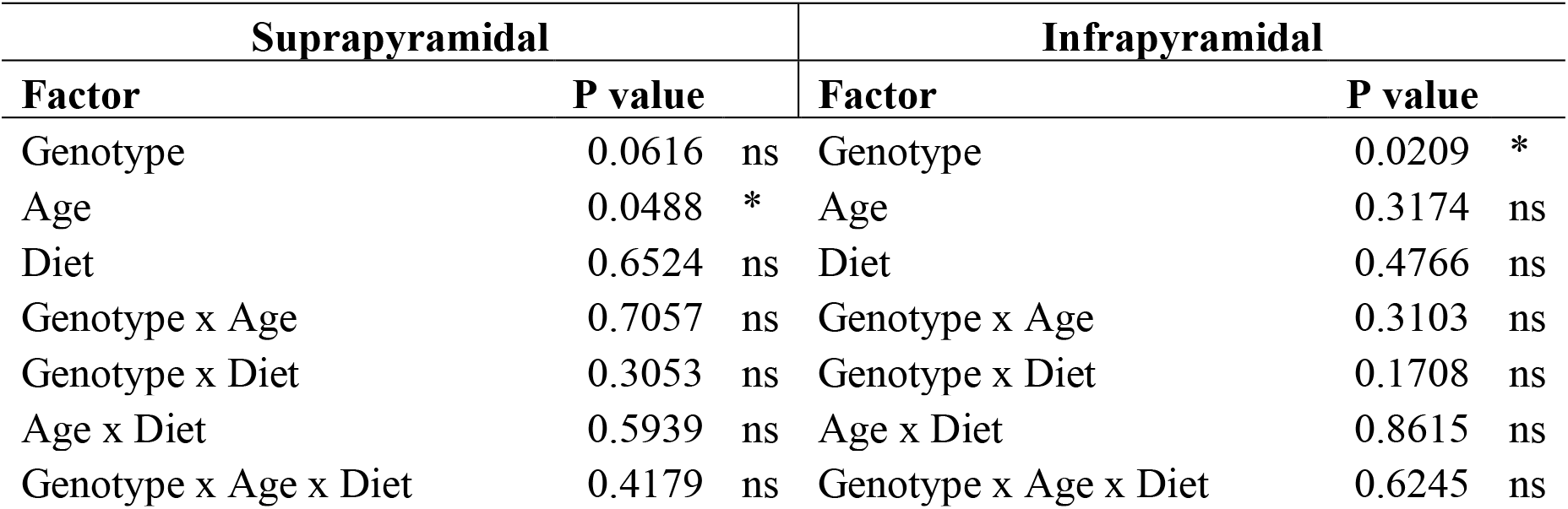
Summary of three-way ANOVA statistical analyses of new-born neurones (BrdU^+^/NeuN^+^) in the suprapyramidal and infrapyramidal blades of the caudal dentate gyrus (DG).

**Table S7.** Summary of repeated measures three-way ANOVA statistical analyses of immature neurone (DCX+) morphology (counts/DG section) in adolescent and adult mice

| Adolescent |  |  | Adult |  |  |
| --- | --- | --- | --- | --- | --- |
| Factor | P value |  | Factor | P value |  |
| Morphology | <0.0001 | **** | Morphology | <0.0001 | **** |
| Genotype | 0.9222 | ns | Genotype | 0.9708 | ns |
| Diet | 0.0729 | ns | Diet | 0.5287 | ns |
| Morphology x Genotype | 0.7891 | ns | Morphology x Genotype | 0.5238 | ns |
| Morphology x Diet | 0.0387 | * | Morphology x Diet | 0.2043 | ns |
| Genotype x Diet | 0.7564 | ns | Genotype x Diet | 0.2269 | ns |
| Morphology x Genotype x Diet | 0.7588 | ns | Morphology x Genotype x Diet | 0.2646 | ns |

**Table S8.**
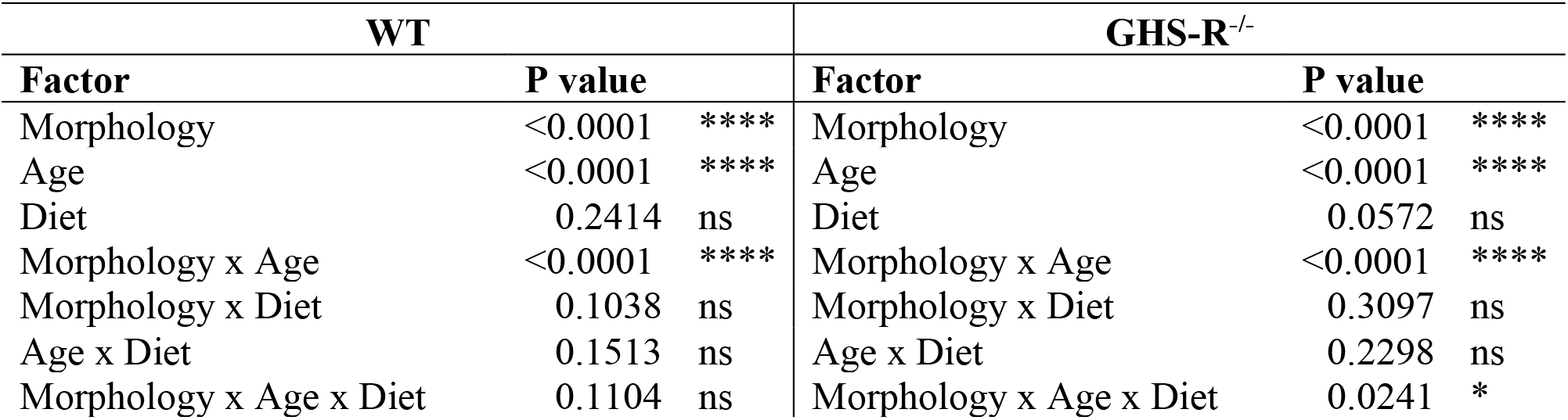
Summary of repeated measures three-way ANOVA statistical analyses of immature neurone (DCX+) morphology (counts/DG section) in WT and GHS-R^−/−^ mice

| WT |  |  | GHS-R<sup>-/-</sup> |  |  |
| --- | --- | --- | --- | --- | --- |
| Factor | P value |  | Factor | P value |  |
| Morphology | <0.0001 | **** | Morphology | <0.0001 | **** |
| Age | <0.0001 | **** | Age | <0.0001 | **** |
| Diet | 0.2414 | ns | Diet | 0.0572 | ns |
| Morphology x Age | <0.0001 | **** | Morphology x Age | <0.0001 | **** |
| Morphology x Diet | 0.1038 | ns | Morphology x Diet | 0.3097 | ns |
| Age x Diet | 0.1513 | ns | Age x Diet | 0.2298 | ns |
| Morphology x Age x Diet | 0.1104 | ns | Morphology x Age x Diet | 0.0241 | * |

**Table S9.**
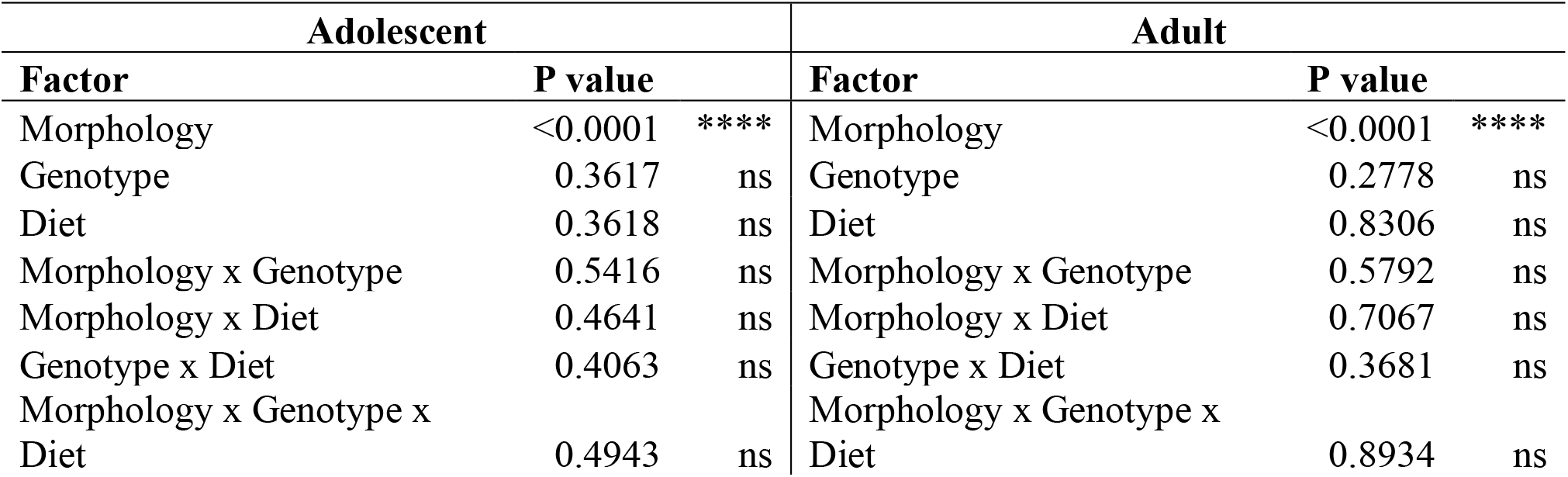
Summary of repeated measures three-way ANOVA statistical analyses of immature neurone (DCX+) morphology (% of total DCX+ cells) in adolescent and adult mice

| Adolescent |  |  | Adult |  |  |
| --- | --- | --- | --- | --- | --- |
| Factor | P value |  | Factor | P value |  |
| Morphology | <0.0001 | **** | Morphology | <0.0001 | **** |
| Genotype | 0.3617 | ns | Genotype | 0.2778 | ns |
| Diet | 0.3618 | ns | Diet | 0.8306 | ns |
| Morphology x Genotype | 0.5416 | ns | Morphology x Genotype | 0.5792 | ns |
| Morphology x Diet | 0.4641 | ns | Morphology x Diet | 0.7067 | ns |
| Genotype x Diet | 0.4063 | ns | Genotype x Diet | 0.3681 | ns |
| Morphology x Genotype x Diet | 0.4943 | ns | Morphology x Genotype x Diet | 0.8934 | ns |

**Table S10.**
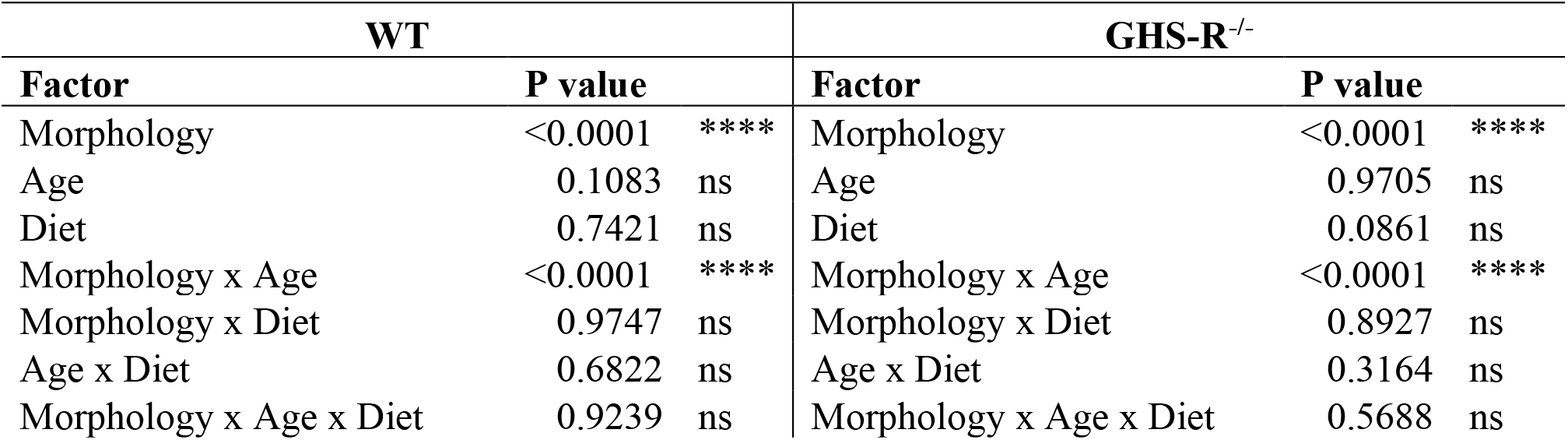
Summary of repeated measures three-way ANOVA statistical analyses of immature neurone (DCX+) morphology (counts/DG section) in WT and GHS-R^−/−^ mice

| WT |  |  | GHS-R<sup>-/-</sup> |  |  |
| --- | --- | --- | --- | --- | --- |
| Factor | P value |  | Factor | P value |  |
| Morphology | <0.0001 | **** | Morphology | <0.0001 | **** |
| Age | 0.1083 | ns | Age | 0.9705 | ns |
| Diet | 0.7421 | ns | Diet | 0.0861 | ns |
| Morphology x Age | <0.0001 | **** | Morphology x Age | <0.0001 | **** |
| Morphology x Diet | 0.9747 | ns | Morphology x Diet | 0.8927 | ns |
| Age x Diet | 0.6822 | ns | Age x Diet | 0.3164 | ns |
| Morphology x Age x Diet | 0.9239 | ns | Morphology x Age x Diet | 0.5688 | ns |

**Table S11.** New neural stem cells (BrdU^+^-Sox2^+^-S100β^−^) in the rostral and caudal poles of the hippocampal dentate gyrus (DG). Data expressed as mean counts (± SEM) per DG section.

|  | WT: AL | GHS-R<sup>-/-</sup> : AL | WT: 5:2 | GHS-R<sup>-/-</sup> : 5:2 |
| --- | --- | --- | --- | --- |
| **Adolescent** |  |  |  |  |
| **Rostral** | 0.25 ± 0.075 | 0.60 ± 0.12 | 0.18 ± 0.075 | 0.38 ± 0.095 |
| **Caudal** | 0.28 ± 0.10 | 0.69 ± 0.11 | 0.18 ± 0.065 | 0.52 ± 0.14 |
| **Adult** |  |  |  |  |
| **Rostral** | 0.10 ± 0.048 | 0.12 ± 0.048 | 0.34 ± 0.14 | 0.15 ± 0.072 |
| **Caudal** | 0.42 ± 0.12 | 0.20 ± 0.061 | 0.18 ± 0.068 | 0.29 ± 0.11 |

**Table S12.** Summary of three-way ANOVA statistical analyses of new neural stem cells (BrdU^+^-Sox2^+^-S100β^−^) in the rostral and caudal poles of the hippocampal dentate gyrus (DG).

| **Rostral** |  |  | **Caudal** |  |  |
| --- | --- | --- | --- | --- | --- |
| **Factor** | **P value** |  | **Factor** | **P value** |  |
| Genotype | 0.1387 | ns | Genotype | 0.0287 | * |
| Age | 0.0075 | ** | Age | 0.053 | ns |
| Diet | 0.929 | ns | Diet | 0.1584 | ns |
| Genotype x Age | 0.0047 | ** | Genotype x Age | 0.0038 | ** |
| Genotype x Diet | 0.168 | ns | Genotype x Diet | 0.3936 | ns |
| Age x Diet | 0.0243 | * | Age x Diet | 0.6713 | ns |
| Genotype x Age x Diet | 0.8305 | ns | Genotype x Age x Diet | 0.1824 | ns |

**Table S13.**
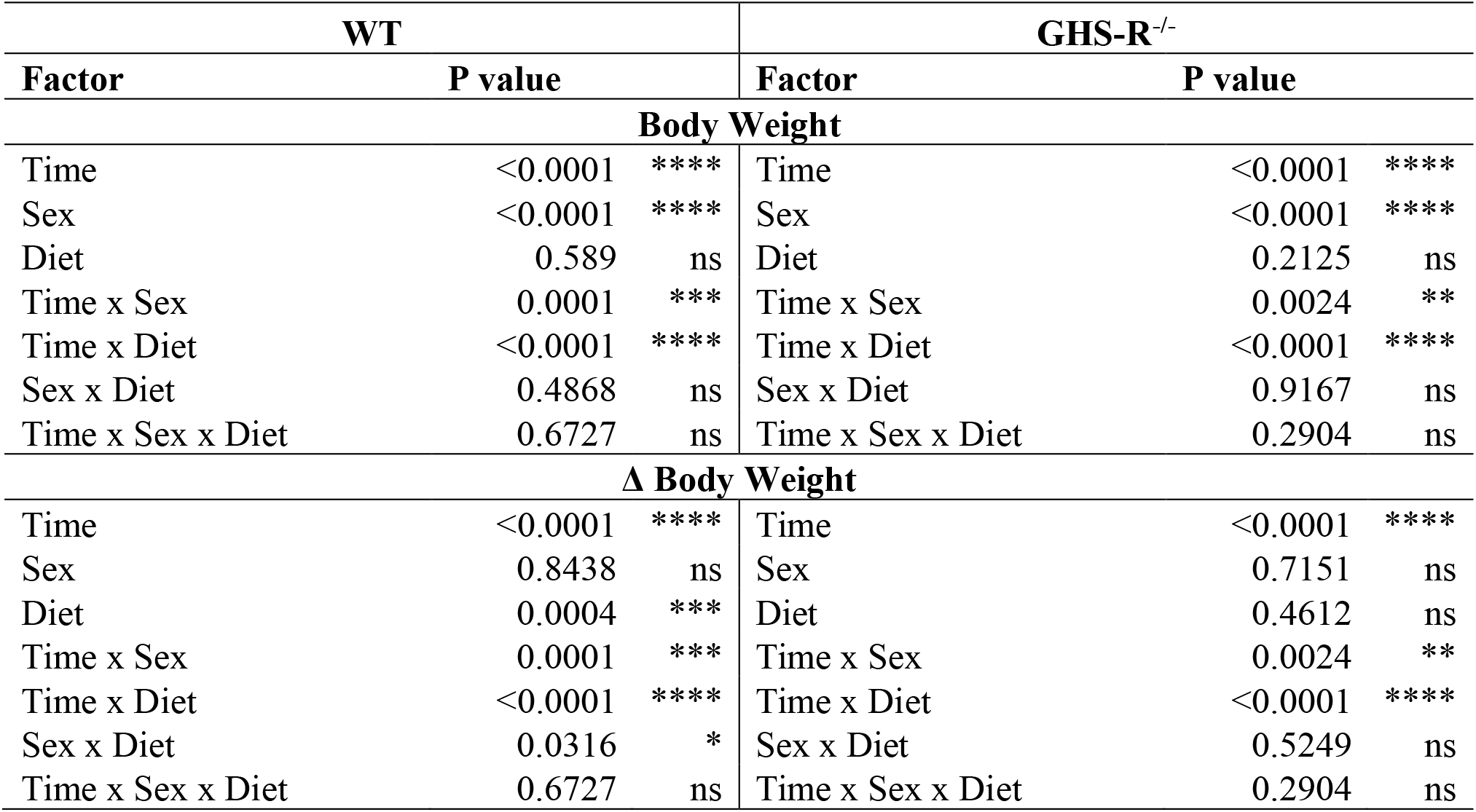
Summary of repeated measures three-way ANOVA statistical analyses of body weight over time and body weight change over time, in male and female, adolescent WT and GHS-R^−/−^ mice

| WT |  |  | GHS-R<sup>-/-</sup> |  |  |
| --- | --- | --- | --- | --- | --- |
| Factor | P value |  | Factor | P value |  |
| Body Weight |  |  |  |  |  |
| Time | <0.0001 | **** | Time | <0.0001 | **** |
| Sex | <0.0001 | **** | Sex | <0.0001 | **** |
| Diet | 0.589 | ns | Diet | 0.2125 | ns |
| Time x Sex | 0.0001 | *** | Time x Sex | 0.0024 | ** |
| Time x Diet | <0.0001 | **** | Time x Diet | <0.0001 | **** |
| Sex x Diet | 0.4868 | ns | Sex x Diet | 0.9167 | ns |
| Time x Sex x Diet | 0.6727 | ns | Time x Sex x Diet | 0.2904 | ns |
| Δ Body Weight |  |  |  |  |  |
| Time | <0.0001 | **** | Time | <0.0001 | **** |
| Sex | 0.8438 | ns | Sex | 0.7151 | ns |
| Diet | 0.0004 | *** | Diet | 0.4612 | ns |
| Time x Sex | 0.0001 | *** | Time x Sex | 0.0024 | ** |
| Time x Diet | <0.0001 | **** | Time x Diet | <0.0001 | **** |
| Sex x Diet | 0.0316 | * | Sex x Diet | 0.5249 | ns |
| Time x Sex x Diet | 0.6727 | ns | Time x Sex x Diet | 0.2904 | ns |

**Table S14.** Summary of repeated measures three-way ANOVA statistical analyses of body weight over time and body weight change over time, in male and female, adult WT and GHS-R^−/−^ mice

| WT |  |  | GHS-R<sup>-/-</sup> |  |  |
| --- | --- | --- | --- | --- | --- |
| Factor | P value |  | Factor | P value |  |
| Body Weight |  |  |  |  |  |
| Time | <0.0001 | **** | Time | <0.0001 | **** |
| Sex | <0.0001 | **** | Sex | <0.0001 | **** |
| Diet | 0.616 | ns | Diet | 0.1405 | ns |
| Time x Sex | 0.0003 | *** | Time x Sex | <0.0001 | **** |
| Time x Diet | <0.0001 | **** | Time x Diet | <0.0001 | **** |
| Sex x Diet | 0.625 | ns | Sex x Diet | 0.5538 | ns |
| Time x Sex x Diet | 0.0439 | * | Time x Sex x Diet | <0.0001 | **** |
| Δ Body Weight |  |  |  |  |  |
| Time | <0.0001 | **** | Time | <0.0001 | **** |
| Sex | 0.0135 | * | Sex | 0.0112 | * |
| Diet | 0.0021 | ** | Diet | 0.0053 | ** |
| Time x Sex | 0.0003 | *** | Time x Sex | <0.0001 | **** |
| Time x Diet | <0.0001 | **** | Time x Diet | <0.0001 | **** |
| Sex x Diet | 0.1399 | ns | Sex x Diet | 0.0482 | * |
| Time x Sex x Diet | 0.0439 | * | Time x Sex x Diet | <0.0001 | **** |

